# 4D Crystallography Captures Transient IF1-Ribosome Dynamics in Translation Initiation

**DOI:** 10.1101/2023.10.27.564398

**Authors:** Ilkin Yapici, E. Han Dao, Shun Yokoi, Ebru Destan, Esra Ayan, Alaleh Shafei, Fatma Betul Ertem, Cahine Kulakman, Merve Yilmaz, Bilge Tosun, Halilibrahim Ciftci, Abdullah Kepceoglu, Jerome Johnson, Omur Guven, Ali Ergul, Brandon Hayes, Yashas Rao, Christopher Kupitz, Frederic P. Poitevin, Mengling Liang, Mark S. Hunter, Pohl Milon, Ayori Mitsutake, Raymond G. Sierra, Soichi Wakatsuki, Hasan DeMirci

## Abstract

Initiation factor 1 (IF1) is one of multiple key ligands involved in the initiation of mRNA translation, a highly dynamic and carefully-orchestrated process. However, details surrounding IF1 transient interactions with the small 30S ribosomal subunit remain incompletely understood despite characterization of unbound and fully-bound 30S states. Improvements in X-ray light sources and crystallographic techniques are now enabling time-resolved structural studies at near-physiological temperature and near-atomic resolution and thus the structural investigation of such dynamic processes. Here, we employed time-resolved serial femtosecond X-ray crystallography (TR-SFX) to probe the binding of IF1 to the small 30S ribosomal subunit in real time. Our time-resolved structural data demonstrates transient cryptic short-, mid-, and long-range allostery among different regions of the small 30 ribosomal subunit during IF1 binding, revealing small- and large-scale protein-target interactions and dynamics within intermediate macromolecular states at unprecedented temporal and spatial resolution. These data represent one of the first such 4D crystallographic studies assessing protein-protein and protein-RNA interactions and could serve as the basis for subsequent studies of the ribosome and of the multitudinous dynamic processes which underpin biology, and therefore, of life.

## Main

Biology is shaped by the dynamic interplay of protein-protein and protein-nucleic acid interactions. Ribosomes are essentially dynamic supramacromolecular machines that orchestrate mRNA translation into protein synthesis (Wimberly et al., 2000; Ramakrishnan, 2002). The four phases of translation comprise initiation, elongation, termination, and recycling (Rodnina, 2018). Initiation is the most highly regulated step and the small ribosomal subunit 30S is known to engage in a highly-orchestrated, multi-layered kinetic process involving initiator tRNA, mRNA and the three initiation factors required for the assembly of the initiation complex (Gualerzi & Pon, 2015; Milon et al., 2010; Milon & Rodnina, 2012;). The binding of the 30S initiation complex to the large 50S subunit induces a series of conformational changes modulating the number of intersubunit bridges (Liu & Fredrick, 2016 ; Yusupov et al., 2001). Initiation factor 1 (IF1) is one of these bridge modulator and required for the fidelity of translation initiation (Milon et al., 2008; Moazed et al. 1995; Qin & Fredrick, 2009).

IF1 binding to the 30S A-site represents an essential step during initiation (Dahlquist & Puglisi, 2000; Zucker& Hershey, 1986). IF1 is critical for cell viability and is an oligomer-binding (OB) fold protein characterized by a five-stranded beta-barrel (Sette et al., 1997). During translational initiation, IF1 kinetically stimulates the 30S initiation complex formation and aids in the dissociation of vacant and mRNA-bound 70S ribosomes (Gualerzi & Giuliodori, 2021; Milon et al., 2012; Pavlov et al., 2008). This smallest initiation factor establishes intimate interactions within the decoding center of 30S which is surrounded by helix 44 (h44), 530 loop, and ribosomal protein uS12 (Carter et al., 2001; Dahlquist & Puglisi, 2000). This binding event involves the flipping of the crucial decoding center residues A1492 and A1493 from h44 of the 16S rRNA. It also induces intertwined mid-scale allostery which is combined with global scale conformational changes (Carter et al., 2001). The rapid IF1 binding and subsequent 30S conformational rearrangements has precluded structural evaluation of initiation complexes and their convoluted dynamics by traditional synchrotron light sources due to radiation-associated structural damage (Garman, 2010; Milon et al. 2012; Sanishvili et al., 2011). Cryogenic temperature restriction has also limited structural observation and analysis of binding and transition-state intermediate complexes. Understanding the *modus operandi* of IF1 in re-engineering the 30S initiation complex conformational landscape to ensure efficiency and accuracy demands advanced, time-resolved structural studies at the atomic scale.

The advent of X-ray free-electron lasers (XFELs) started a new era in structural biology, introducing an unparalleled capability to visualize transient states of biomolecules with ultra-short, high-intensity X-ray pulses (Chapman et al., 2011; Pellegrini, 2020). Methods developed for macromolecular studies at XFELs employ a different data acquisition compared to synchrotrons, commonly termed “diffract-before-destroy” (Neutze et al., 2000). Rather than exposing one or few monolithic large macromolecular crystals to obtain a complete data-set, the ultrafast pulses diffract numerous nano- and microcrystals in succession immediately before destroying them (Chapman et al., 2011). Serial data acquisition using fresh crystals minimizes the problem of radiation damage and has enabled the determination of crystal structures to high resolution at non-cryogenic temperatures (Boutet et al., 2012; Chapman et al., 2011; Martin-Garcia et al., 2016; Neutze et al., 2000; Takaba et al., 2023;). Temperature is a critical physical parameter in protein dynamics (Keedy et al., 2015) and also crystallization of biomacromolecules due to its impact on enthalpy balance (Russo Krauss et al., 2013). Biomacromolecular structures are sensitive to temperature fluctuations, particularly in the case of nucleic acids which contain fewer hydrogen bonds compared to proteins (Liao et al., 2021; Ryals et al., 1982; Vogt & Argos, 1997). Therefore, determining protein and nucleic acid complex structures at ambient temperature is crucial for improved characterization of their function and dynamics. Multiple sample delivery approaches have been developed and described for application at XFELs. One of these approaches is the concentric-flow microfluidic electrokinetic sample holder (coMESH) (Fig. 1a), previously employed for the investigation of megadalton-sized, fragile crystal structures at near-physiological temperatures (Dao et al., 2018; O’Sullivan et al., 2018; Sierra et al., 2016).

**Fig.1.**
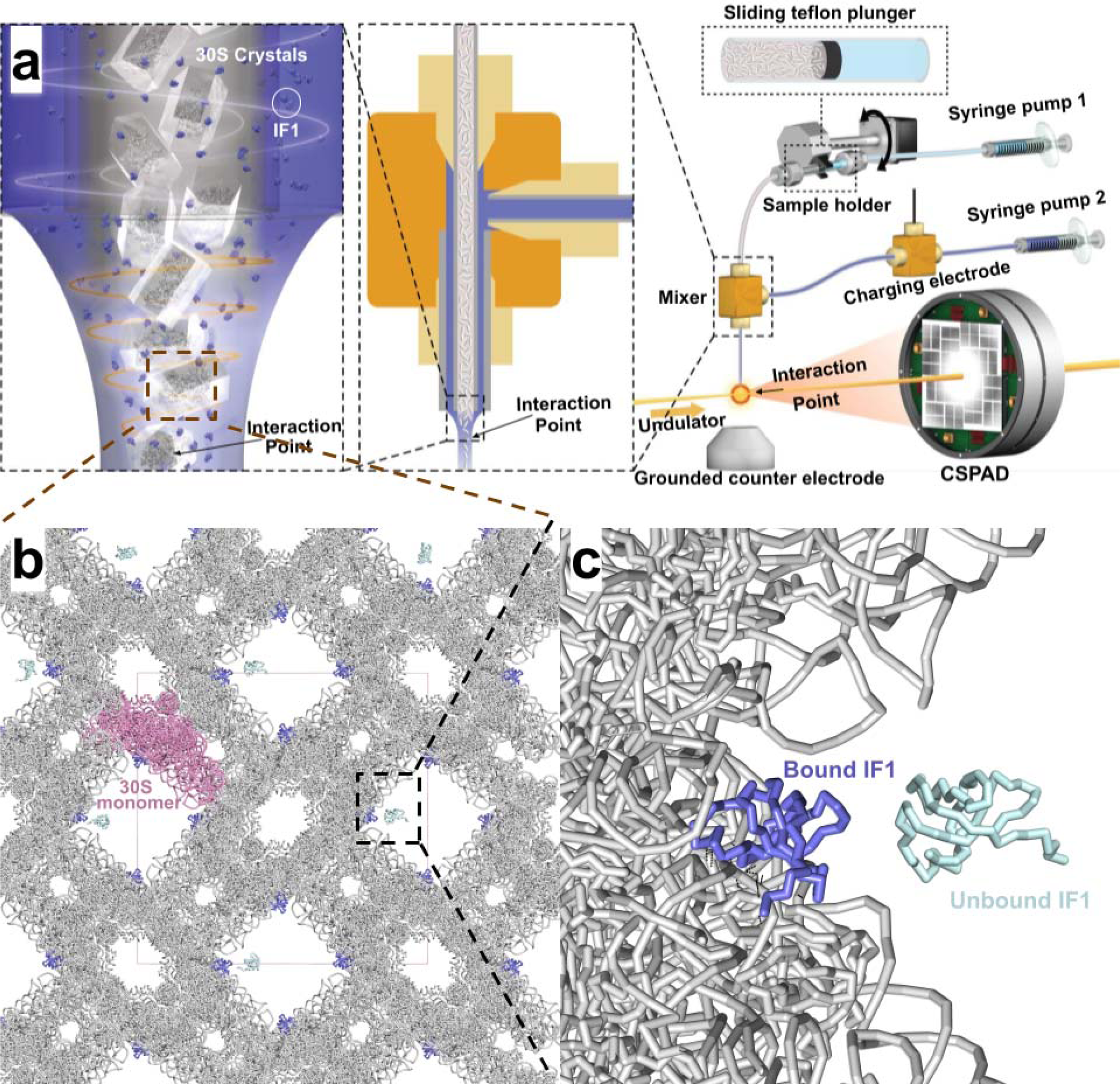
Illustration of the concentric-flow microfluidic electrokinetic sample holder (co-MESH), 30S and IF1 crystal packing. **a, (right)** >A continuous inner capillary (100 μm × 160 μm × 1.5 m) facilitated the flow of a liquid jet, which included 30S microcrystals and their mother liquor (16% v/v MPD; ; colored in gray). Electro-focusing of the liquid jet was achieved by charging a sister liquor containing IF1 solution (colored in purple) using a high-voltage power supply. Teflon plunger (colored in black) separated the sample reservoir from the driving fluid (colored in light blue) . The reservoir was mounted on an anti-settling device, which rotated at an angle about the capillary axis to keep the protein crystals homogeneously suspended in the mother liquor. **(middle)** A mixer (indicated within the dashed black rectangle) connected the two capillaries (colored in gray) in a concentric manner.The interaction between the liquid jet and the LCLS pulses occurred at the point marked by the orange circle. **(left)** Schematic representation of electro-spinning and focusing and mixing of co-flowing 30S microcrystals with IF1 solution. **b,** The packing arrangement of the 30S^TR-SFX^ crystal lattice indicating 170 Å wide solvent channels from one axis. An individual 850 kDa 30S monomer is shown in pink. **c,** A close-up view of the solvent channel and IF1-30S binding interphase is shown. 30S bound IF1 is colored in slate and unbound IF1 colored palecyan and shown only for illustration purpose of the fitting to the solvent channel.

Going beyond static crystal structures, sample delivery methods at XFELs have been extended to pursue the study of transient intermediates of biomacromolecules (Johansson et al., 2017; Khakhulin et al., 2020). Pump-probe and mix-and-inject XFEL serial crystallography strategies have been described, offering new approaches to evaluate protein-protein or protein-RNA interactions as they occur, and thus opening a new area of time-resolved serial femtosecond crystallography (TR-SFX) (Orville, 2018, 2020; Pandey et al., 2020; Stagno et al., 2017). For example, mix-and-inject SFX was used to capture the intermediate state of a small-molecule ligand and its associated riboswitch RNA, revealing local conformational changes around the ligand binding pocket in real time. Other examples include the catalysis of small-molecule ceftriaxone cleavage by the bacterial β-lactamase enzyme BlaC (Olmos et al.,2018) and intermediate states of isocyanide hydratase (ICH) enzyme catalysis during hydrolysis (Dasgupta et al., 2019). Similar structural studies of paired large macromolecular interactions, e.g. protein-protein or protein-RNA interactions, have yet to be described. Ribosome complexes and their crystals are ideal model systems given multiple associated large substrates and their wide-open solvent channels, permitting the diffusion of protein factors such as IF1 to its decoding center binding pocket (Carter et al., 2001) (Fig. 1b,c).

Cryogenic crystal and cryoEM structures of bacterial translation initiation complexes, including the fully-bound state of 30S and IF1 (hereafter referred as 30S^HOLO^), have been previously reported (Carter et al., 2001; Hussain 2016; Myasnikov et al., 2005) (Fig. 2a,b). The mechanism of these initial binding steps of IF1 to the decoding center remains unclear, leaving an incomplete understanding of dynamics of initiation complex formation. Here, mix-and-inject data acquisition of the 8.2 kDa IF1 protein and crystals of the ∼1 MDa 30S was achieved by further adaptation of the coMESH sample-delivery method. Our resulting TR-SFX structure (hereafter referred as 30S^TR-SFX^) captures intermediate states in real time to reveal multiple protein-protein and protein-RNA interactions at 3.59 Å resolution. The transient binding interactions between the IF1 and the decoding center were captured in ∼200 millisecond temporal resolution. Notable changes were observed within critical decoding residues A1492 and A1493 within h44, and the 530 loop region of 16S rRNA, in addition to its interaction with ribosomal protein uS12. In particular, intermediate conformational changes and the uncoupling of h44 motion, and global scale allosteric domain closure from head rotation of IF1 binding are observed. These time-resolved data provide the first snapshot to the developing cinematographic view of the all-important translation initiation process.

**Fig. 2.**
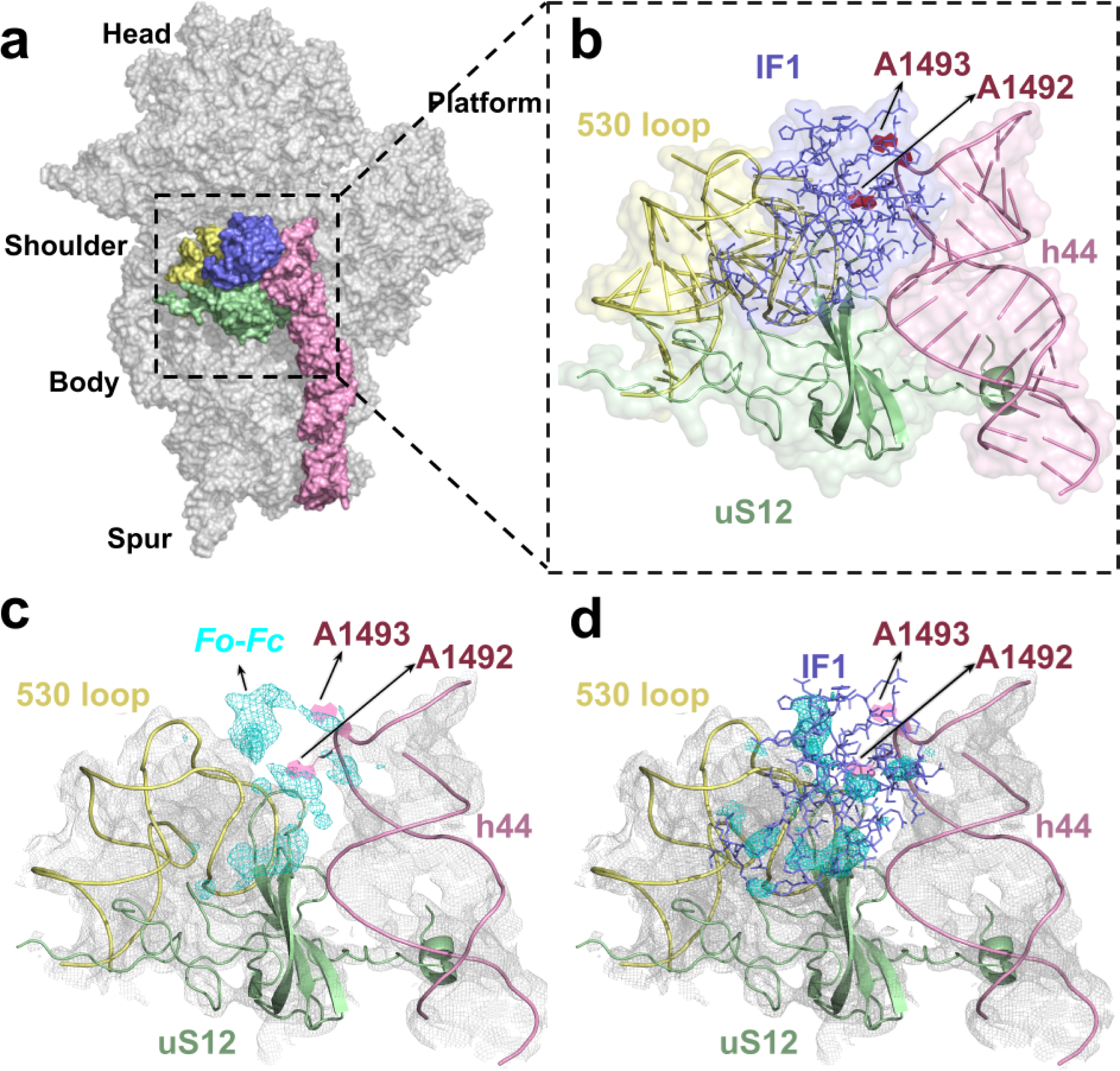
IF1 on its path to binding to the 30S ribosomal subunit. **a,** Surface representation of the “chick view” showing the positions of IF1 colored in slate, h44 colored in pink, ribosomal protein uS12 colored in green, 530 loop colored in yellow. **b,** Close up view of the inset in panel a, cartoon representation of IF1 binding pocket, the decoding residues A1492 and A1493 colored in red and represented in filled cartoons. **c,** Unbiased *Fo-Fc* omit electron density map without an IF1 model, colored in light blue and contoured at 3σ level; *2Fo-Fc* electron density map of 30S colored in cyan and contoured at 1σ level. **d,** *2Fo-Fc* electron density map after refinements with IF1, contoured at 1σ level.

### 30S-IF1 complex formation captured by TR-SFX

Our TR-SFX method based on coMESH injection technology extends beyond previously-described implementations to facilitate the active mixing of 30S microrycstals and IF1 solution through electrospinning. Separate solutions of *Thermus thermophilus* (*T. thermophilus*) 30S microcrystals and purified IF1 were loaded into separate, concentric capillary tubing, with the tubing arranged to mix the solutions immediately prior to the XFEL beam (Fig. 1a), relying on different charge densities of each solution to achieve mixing under applied voltage. Flow rates in each capillary measured 1 ul/min resulting in an associated flow velocity of 0.212 cm/s. When an applied voltage is introduced, the jet’s velocity falls within the range of 2.12 to 6.36 cm/s corresponding to the timescale of interactions spanning from 80 to 240 milliseconds. A complete dataset acquisition was achieved using 300 microliters of ribosome microcrystal slurry delivered to the Coherent X-ray Imaging (CXI) Instrument at the Linac Coherent Light Source (LCLS) at a scheduled beamtime (cxils9717). The full time-resolved diffraction dataset after ∼200 milliseconds mixing for 30S-IF1 complex structure (30S^TR-SFX^) containing both 850 kDa 30S, and also partially occupied 8.2 kDa IF1 was collected from microcrystals in the P4_1_2_1_2 space group in 216 minutes and 58 seconds. The initial rigid body refinement against the apo 30S structure (30S^APO^)(PDB ID: 4DR1) resulted in a refined model with a *R_work_/R_free_*of 0.26/0.29 (Fig. 2c). The IF1 was omitted during the initial refinement; however, the corresponding unbiased *Fo-Fc* electron density maps unambiguously reveal the partial occupancy of the incoming IF1 to the binding cavity formed between h44 and 530 loop region of the 16S rRNA and uS12 protein of 30S (Fig. 2c). Subsequently, the IF1 model was rebuilt manually by using the crystal structure of 30S^HOLO^ (PDB ID: 1HR0) as a guide model by using *COOT* (Carter et al., 2001; Emsley & Cowtan, 2004) (Fig. 2d). The resolution cut-off is 3.59 Å with 100 % completeness, and after 19 rounds of iterative refinement and model building, the final structure resulted in *R_work_/R_free_*of 0.23/0.28. The statistical information for the data is available in Extended Data Table 1.

### Transient IF1 interactions with h44, 530 loop and protein uS12

The 30S^TR-SFX^ structure explores both the protein-nucleic acid and protein-protein interaction dynamics. Extended Data Fig. 1 reveals the electrostatic surface charge of IF1 and its surroundings on the 30S. Mostly neutral charged IF1 binds within the cleft formed by the h44, 530 loop and the positively charged ribosomal protein uS12 near the decoding center, suggesting the potential role of uS12 in steering IF1 toward h44 which corroborates with results of molecular dynamics (MD) simulation (Fig. 4). Furthermore, IF1’s negatively charged regions interact with the uS12 protein while its positively charged regions interact with the h44 and 530 loops via their negatively charged phosphate backbone. Stabilization of IF1 on its binding site is a dynamic process, ending with certain bonds preserving the bound state. Besides unraveling IF1’s mode of binding during preinitiation, our study showed the unprecedented atomic detailed view of the initial binding process of IF1 (Fig. 4).

### Allostery at the decoding center

To reveal the structural dynamics of the decoding center during different stages of IF1 binding, we compared and analyzed the experimental electron density maps from three ribosome crystal structures of: i) 30S^APO^ (PDB ID: 4DR1), ii) 30S^TR-SFX^ (PDB ID: 8WRC) and iii) 30S^HOLO^ (PDB ID: 1HR0) (Fig. 3) (Carter et al., 2001; DeMirci et al., 2013). This comparison revealed previously unobserved and cryptic intermediate structural information of h44 regions engaging in IF1 binding (Fig. 3). The superposition of the three structures confirmed the remarkable flexibility of universally-conserved decoding residues A1492 and A1493 during the initial stages of IF1 binding (Fig. 3; Supplementary Video 1). Based on experimental time-resolved 2*Fo-Fc* electron density map collected at ∼200 milliseconds, the resulting 30S^TR-SFX^ intermediate structure with IF1 on its path to 30S decoding center has weaker electron density for A1493 and better defined density for A1492. Although the 30S^HOLO^ structure has a well-defined electron density for residues A1492 and A1493, the 30S^APO^ structure shows these residues in a tucked-in conformation with a less well-defined electron density. (Fig. 3b,g). These results suggest that initial IF1 binding steps capture alternative local conformational changes on h44 of 16S rRNA. Extensive one microsecond molecular dynamic (MD) simulations performed for the dissociation of IF1 from 30S revealed novel intermediate conformation of decoding residues (see below). Further, it may provide computational details of the choreography and how IF1 dissociation leads to transitioning conformational changes in the decoding center.

**Fig. 3.**
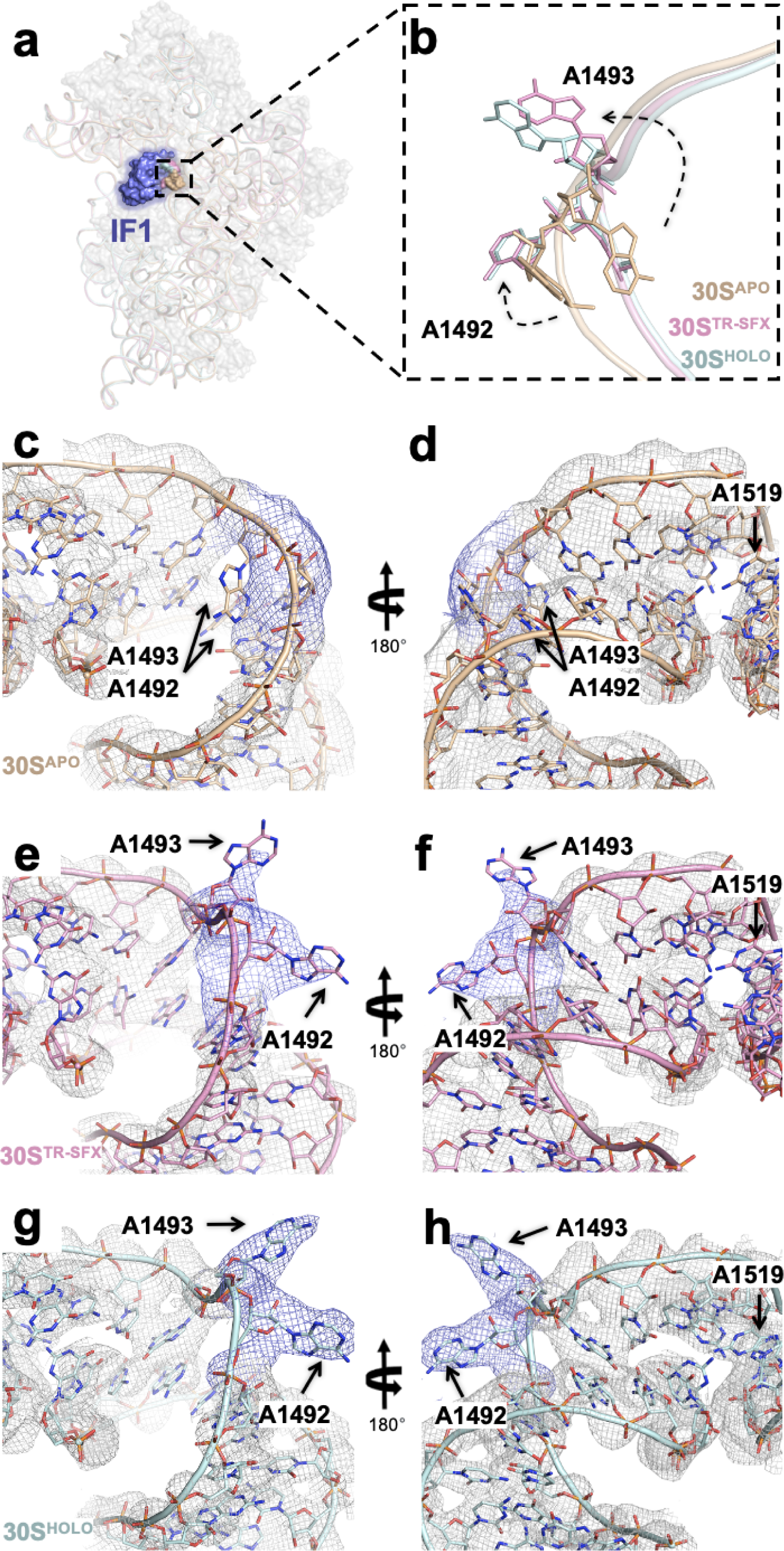
Conformational changes of decoding residues A1492 and A1493 during IF1 binding to 30S. **a,** Chick view of the 30S ribosomal subunit **b,** 30S^TR-SFX^ intermediate state structure is superposed with 30S^APO^ and 30S^HOLO^ structures (PDB ID: 4DR1 & 1HR0, respectively) to represent the plasticity of the decoding residues A1492 and A1493. **c-d,** The 30S^APO^ structure is colored in wheat and *2Fo-Fc* electron density map is colored in gray and location of the decoding residues A1492 and A1493 are indicated by arrows. *2Fo-Fc* electron density maps are contoured at the 1σ level (colored contour levels remains same for all the panels). **e-f,** the 30S^TR-SFX^ intermediate state structure is colored in pink while **g-h,** 30S^HOLO^ colored in palecyan.

**Fig. 4.**
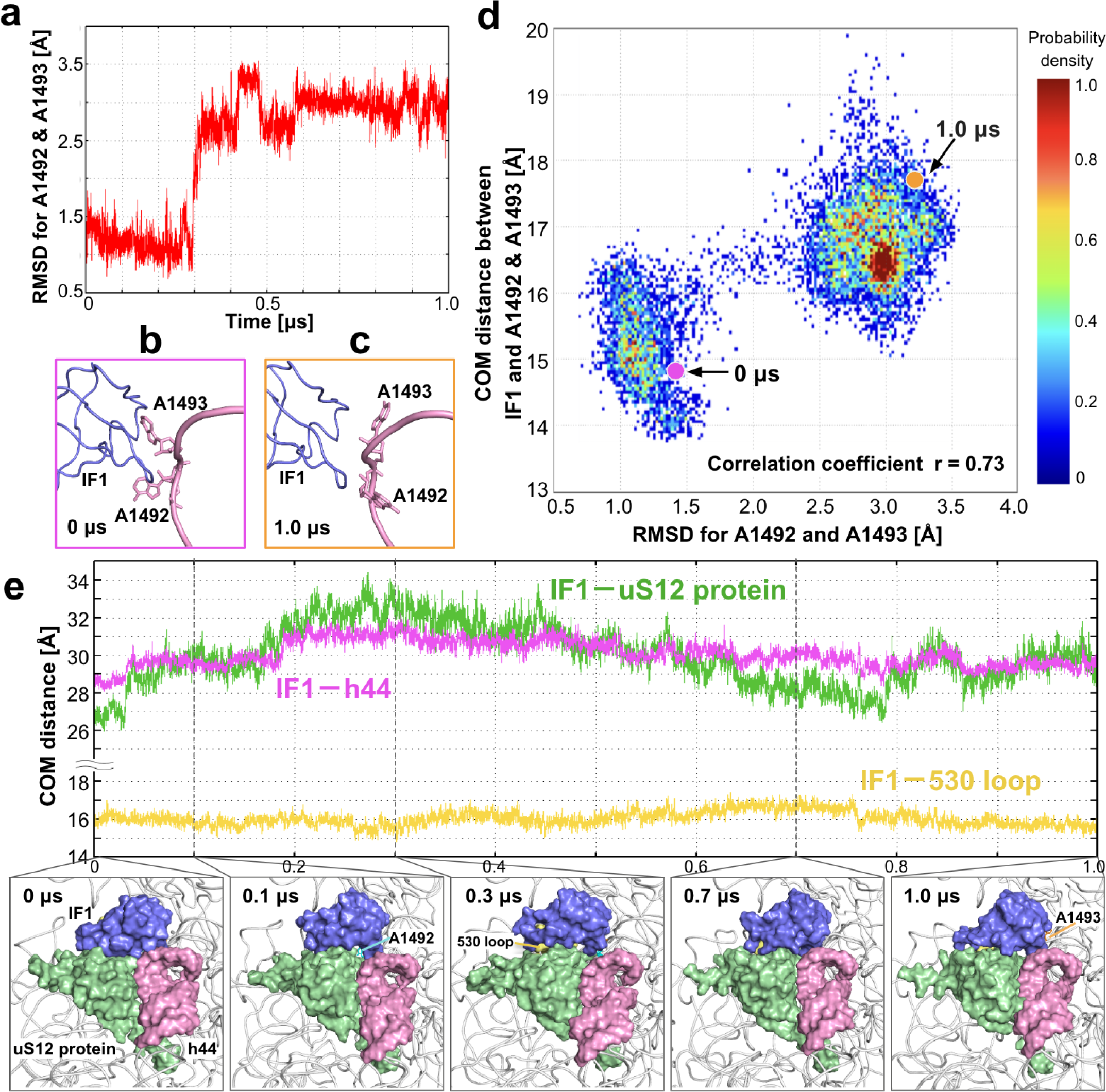
The dynamics of the decoding center in MD simulation of IF1 bound 30S. **a,** Time series of RMSD for A1492 and A1493. RMSD was calculated for all atoms except hydrogen atoms in A1492 and A1493, using the 30S^TR-SFX^ structure as a reference structure.The RMSD changed significantly around 300 ns, when the side chain of A1492 transitioned into the tucked-in conformation. **b,** Correlations between RMSD for A1492-A1493 and the distance between centers of mass (COM distance) of IF1 and A1492-A1493. The COM distance was calculated using all atoms except hydrogen atoms in the entire IF1 structure and A1492-A1493, respectively. There was a strong correlation between the RMSD and the COM distance, with a correlation coefficient of 0.73. The values at 0 μs and 1 μs correspond to the pink and orange dots, respectively, and the structures are shown in the bottom. The conformational changes in A1492 and A1493 led IF1 to move away from the decoding center. **c,** Time series of COM distances of IF1 and uS12 protein, of IF1 and h44, and of IF1 and 530 loop. IF1, uS12 protein, h44 (near A1492 and A1493), and 530 loop are colored in blue, green, pink, and yellow, respectively. The several characteristic structures over the time series are also shown. IF1, uS12 protein, and h44 were packing at 0 ns, and the interactions between them were strong. Their interactions were loosened by 300 ns, and IF1 dissociated a little from the decoding center. After the interaction between IF1-uS12 protein and IF1-h44 became loose, the COM distances of IF1 and uS12 protein and of IF1 and h44 were significantly fluctuating. On the other hand, the COM distance of IF1 and 530 loop was stable.

### MD simulation of the 30S-IF1 complex

We performed one microsecond all-atom MD simulation using the 30S^TR-SFX^ structure to investigate the dynamics of the interface interactions between IF1 and the 30S in detail. For investigating local conformational changes of A1492 and A1493, the root mean square deviation (RMSD) for all atoms except hydrogen atoms in A1492 and A1493 from the initial structure as a reference was calculated as shown in Fig. 4a. The conformations of A1492 and A1493 changed significantly during the MD simulation. Around 300 ns, the side chain of A1492 transitioned into the tucked-in conformation, resembling the 30S^APO^ state as shown in the Fig. 4b,c. After this A1492 conformational change, the structural fluctuations of A1493 increased, resulting in the A1493 side chain also swinging away from IF1 (Supplementary Video 2).

In the simulation, we found that IF1 slightly moved away from the decoding center along with these conformational changes. Fig. 4d shows the distance between centers of mass (COM distance) of IF1 and A1492–A1493 as a function of the RMSD. While the COM distance of IF1 and A1492–A1493 was 14.8 Å initially, the average COM distance over the MD simulation was 16.3 ± 0.9 Å, which indicates that the distance was slightly increasing over time. Furthermore, we found that there was a strong correlation between the RMSD of A1492–A1493 and the COM distance between IF1 and A1492–A1493, with a correlation coefficient of 0.73. While the average COM distance was 15.2 ± 0.6 Å when the RMSD was around 1 Å, the average distance was 16.8 ± 0.6 Å when the RMSD was around 3 Å. Thus, the structural change to the tucked-in conformations of A1492–A1493 caused IF1 to dissociate and fluctuated significantly in the decoding center (Fig. 4d, Supplementary Video 2). Six additional 100 ns MD simulations using the structure at 1 μs as the initial structure were performed, and we found that A1493 formed the fully tucked-in conformation, which is similar to the conformation in 30S^APO^ state (Supplementary Video 3). From the MD simulation, the structural fluctuation of A1493 was large, which is consistent with the experimental results of weak electron density of A1493 in the 30S^TR-SFX^ structure.

Interestingly, the conformational change of A1492 and A1493 and the dissociation of IF1 might be caused by the weakening of the interfacial interactions between IF1 and uS12 protein and between IF1 and h44. Fig. 4e shows the COM distances between IF1 and uS12 protein, IF1 and h44, and IF1 and 530 loop respectively. The distances for uS12 protein and h44 were 26.9 Å and 28.7 Å, respectively; while their average distances over the course of the one microsecond MD simulation were 30.0 ± 1.6 Å and 30.1 ± 0.8 Å. At 0 ns, IF1, uS12 protein, and h44 were tightly packing, and the interactions between them were strong. However, their interactions were loosened by 300 ns, and IF1 dissociated slightly from the decoding center. After the interaction between IF1-uS12 protein and IF1-h44 became loose, the COM distances of IF1 and uS12 protein and of IF1 and h44 were significantly fluctuating (Supplementary Video 4). On the other hand, the initial COM distance of IF1 and 530 loop was 15.7 Å, and the average COM distance was 16.0 ± 0.4 Å, indicating that the interaction between them was stable. Therefore, it is suggested that the dissociation of IF1 from the decoding center observed in the MD simulation was attributed primarily to the disengagement of the IF1-uS12 and IF1-h44 interface interactions.

### IF1 stabilized mid-scale allostery in h44

In our 30S^TR-SFX^ complex structure, h44 is captured in significantly altered conformations at two distinct regions of the 16S rRNA (Fig. 5a). We mapped these perturbations of the base interactions within h44 by observing the presence and absence of hydrogen bonds between h44 base pairs using *RNApdbee 2.0* and *VARNA* programs (Antczak et al., 2014 2018; Darty et al., 2009; Zok et al., 2018). The 30S^APO^ structure exhibits hydrogen bonds near the decoding center, specifically between residue pairs C1412-G1488, A1413-G1487, U1414-G1486, G1415-U1485 (Fig. 5b,c) G1423-C1477, C1424-G1476, U1425-G1475, C1426-G1474, and U1427-A1473 (Fig. 5b,d). The 30S^HOLO^ structure lacks interactions between the residue pairs A1413-G1487 and U1425-G1475 (Fig. 5h,i,j). In the 30S^TR-SFX^ structure, we observed that interaction with IF1 stabilizes alternate transient hydrogen bond networks, which causes changes in bond distances between the residue pairs C1412-G1488, A1413-G1487, U1414-G1486 and G1415-U1485 (Extended Data Fig. 2). Furthermore, the hydrogen bond network between the residue pairs G1423-C1477, C1424-G1476, U1425-G1475, C1426-G1474, and U1427-A1473, observed in the 30S^APO^ and 30S^HOLO^ structures, seem to be transiently disrupted during the binding process as observed in our 30S^TR-SFX^ structure (Fig. 5e,f,g). Moreover, by demonstration of the intermediate state through *VARNA*-generated flattened secondary structures we observed the presence of single hydrogen bonds in non-canonical residue pair U1414-G1487, as well as the occurrence of extra double hydrogen bonds between residue pair C1426-G1475 (Fig. 5e,f,g). Our analysis reveals that, even before IF1 fully binds, the factor disrupts the equilibrium of transient molecular energy minima excursions within the h44. At this time point, one of the energetically favorable excursions by the perturbation of this equilibrium by IF1 collision may lead to the transient stabilization of an intermediate state. Consequently, 30S^TR-SFX^ structure serves as the first ribosome structure showing the short-lived and reversible conformational excursions on h44 region of the 16S rRNA during the initial binding state of IF1 to the 30S. The perturbations of the base interactions within h44 were also observed in the one microsecond MD simulation, and the conformations in the MD simulation were similar to the 30S^APO^ state (Supplementary Video 5, Supplementary Video 6, Supplementary Video 7, Supplementary Video 8).

**Fig.5.**
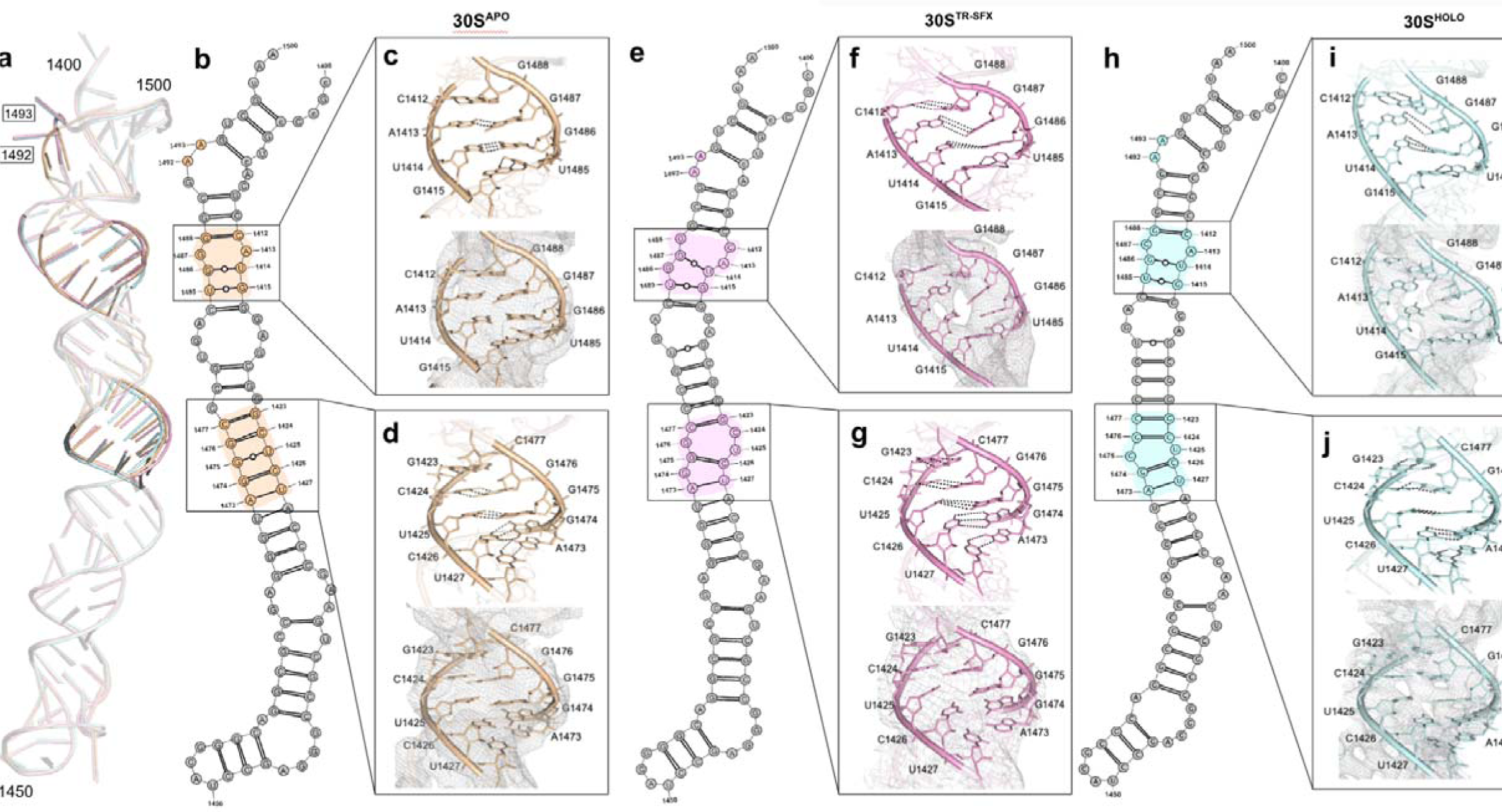
The perturbations of the base interactions observed in the h44 region of the 16S rRNA structure during IF1 binding. The rearrangements of the base interactions are detected by measuring the distances between base pairs and assessing the presence of chemical interactions based on proximity. **a,** 30S^APO^, 30S^TR-SFX^ and 30S^HOLO^ structure of h44 superposed and decoding residues A1492 and A1493 and perturbed residues are colored in wheat, pink and palecyan respectively. **b,** The secondary structure of 30S^APO^ (wheat) h44 region of the 16S rRNA. **c,f,i,** The perturbations of the base interactions in the 30S^APO^, 30S^TR-SFX^ and 30S^HOLO^ h44 structure are depicted, demonstrating the distances between the residue pairs C1412-G1488, A1413-G1487, U1414-G1486, and G1415-U1485 respectively. **e,** The secondary structure of 30S^TR-SFX^ (pink) h44 region of the 16S rRNA. perturbations of the base interactions in the 30S^APO^, 30S^TR-SFX^ and 30S^HOLO^ h44 structure are **d,g,j,** The depicted, demonstrating the distances between the residue pairs G1423-C1477, C1424-G1476, U1425-G1475, C1426-G1474, and U1427-A1473 respectively **h,** The secondary structure of 30S^HOLO^ (palecyan) h44 region of the 16S rRNA.

### TR-SFX decouples mid- and global-scale IF1 allosteric intermediates

In our complete 16S rRNA pairwise comparative analysis, the main chain phosphate positions of the 30S^APO^, 30S^TR-SFX^ and 30S^HOLO^ structures were superposed (Fig. 6). Upon aligning the 30S^APO^ structure with the 30S^TR-SFX^ structure, the maximum spatial shift of 6 Å between residues A1408-G1422 resulted in h44 motions relative to the starting 30S^APO^ structure (Fig. 6a,e,f). Within the 30S^TR-SFX^ structure, the initial partial binding of IF1 led to directed lateral shifts of approximately 2.0 Å in the vicinity of residues G75-U98 and A463-U480 (Fig. 6f). Additionally, a comparative analysis between the 30S^TR-SFX^ and 30S^HOLO^ structures reveals a slight shoulder displacement of up to 2 Å between A453-G485 residues (Fig. 6b,g,h). When 30S^TR-SFX^ and 30S^HOLO^ structures are superposed, the intermediate binding of IF1 to the 30S structure reveals h44 motions and head movements spanning a range of 2.0-4.0 Å between residues A938-G1454 (Fig.6 g,h), opposite to the behavior observed from comparing the 30S^APO^ and 30S^TR-SFX^ structures (Fig. 6f, Supplementary Video 9). However, in contrast to the pairwise superpositions involving all three structures of 30S^APO^&30S^TR-SFX^ or 30S^TR-SFX^&30S^HOLO^, we observed that the 30S^HOLO^ structure leads to significant body movements in local positions when aligning with the 30S^APO^ and 30S^HOLO^ structures. These movements result in notable perturbations in the distances between residues, particularly in the regions spanning from 402 to 502 and in the vicinity of residues 902 to 1502 (Fig. 6c). The 30S^TR-SFX^ data not only help to capture intermediate phases within the IF1-mediated binding process to the 30S structure but also effectively distinguishes and isolates a cryptic intermediate step between two distinct 30S^APO^ (t=0) and 30S^HOLO^ (t=∞) states (Fig. 6d). Collectively, these findings indicate that 30S^TR-SFX^ structure is a novel global intermediate IF1 binding state.

**Fig. 6.**
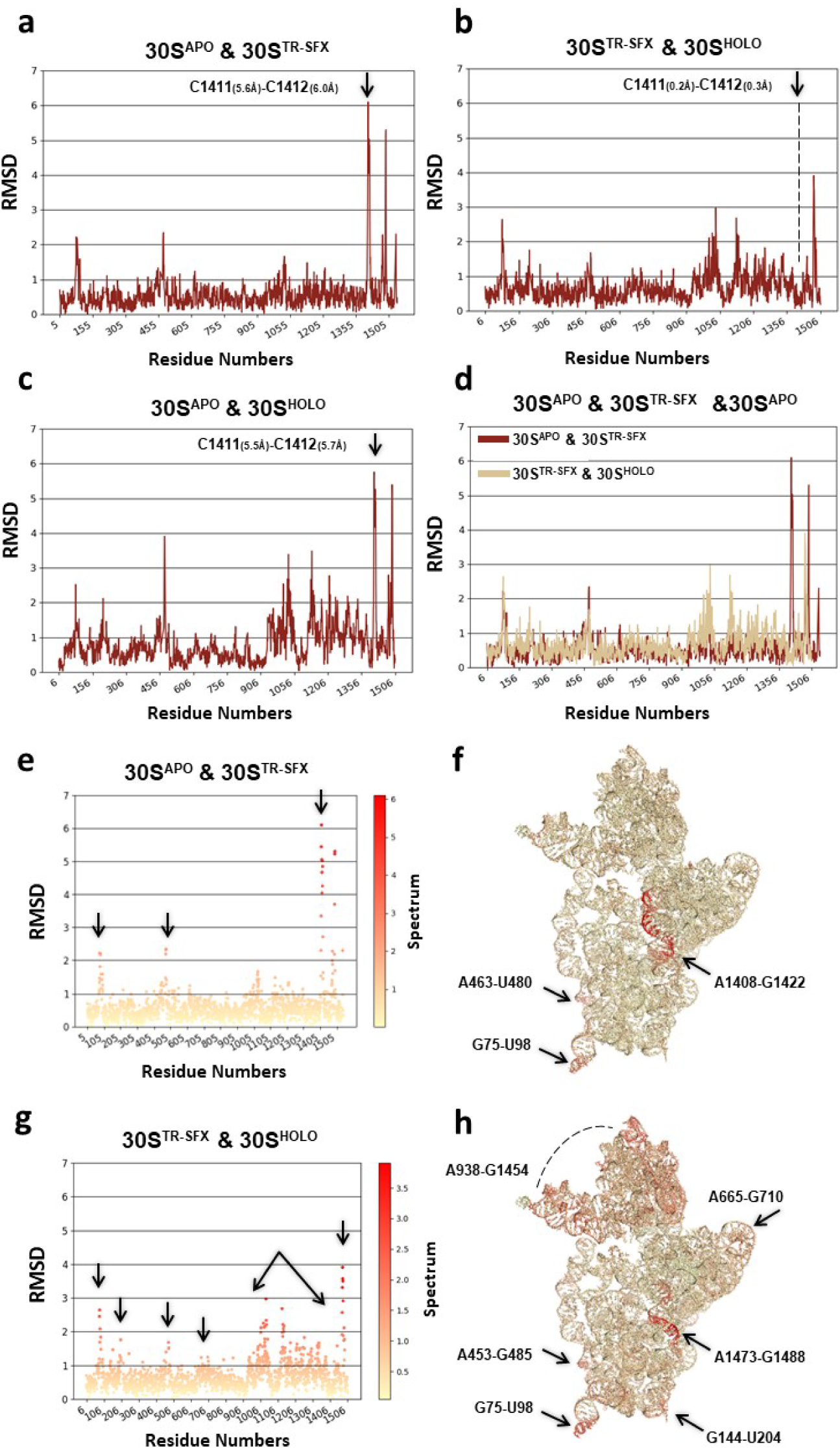
Interatomic distance analysis of 30S^APO^, 30S^TR-SFX^, and 30S^HOLO^ structures. **a,e,f,** Comprehensive analysi of both two-dimensional (2D) and three-dimensional (3D) pairwise distances between the 30S^APO^ and 30S^TR-SF^ structures. The alignment of the 30S^APO^ structure with the 30S^TR-SFX^ structure reveals a significant maximum spatia shift of 6 Å between residues A1408-G1422, which resulted in substantial h44 motions relative to the initial 30S^AP^ structure. The initial binding of IF1 led to lateral shifts of approximately 2.0 Å, particularly in the vicinity of residue G75-U98 and A463-U480. **b,g,h,** An in-depth examination of 2D and 3D pairwise distances between the 30S^TR-SFX^ an 30S^HOLO^ structures reveals a subtle displacement of up to 2-3 Å between residues A453-G485, G144-U204, A665-G710 and G75-U98. Upon superimposing the 30S^TR-SFX^ and 30S^HOLO^ structures, the intermediate binding of IF1 to the 30 structure initiated h44 motions and head movements spanning a range of 2.0-4.0 Å, primarily between residues A938 G1454. **c,** The 2D pairwise distances between the 30S^APO^ and 30S^HOLO^ structures indicates 30S^HOLO^ structure induce substantial repositioning in localized positions when aligned with the 30S^APO^ and 30S^HOLO^ structures. These movement led toperturbations in the distances between residues, particularly within the regions spanning from 402 to 502 and i the proximity of residues 902 to 1502. **d,** Superimpositions involving all three structures of 30S^APO^ & 30S^TR-SFX^ o 30S^TR-SFX^ & 30S^HOLO^.

To investigate large-scale movements of 30S in the MD simulation, the root-mean-square fluctuation (RMSF) from the average structure is shown in Extended Data Fig. 3. There are large structural fluctuations around G75-U98, C150-G168, A453-G485, G674-G714, and A938-G1454 (specifically, C1254-C1284), which correspond to large movements as results of principal component analysis (Supplementary Video 10, Supplementary Video 11). The large-scale movements of 30S in the MD simulation had similar structural fluctuation tendency as those in the 16S rRNA pairwise comparative analysis of 30S^TR-SFX^ and 30S^HOLO^ except for C150-G168, G674-G714, C1254-C1284 and the decoding center of h44. The similarity may be caused by ambient temperature. The different results for C150-G168, G674-G714, and C1254-C1284 may be caused by the effect of crystal packing. While the outside regions of 30S fluctuated, the decoding center did not fluctuate significantly in the MD simulation. Even though the decoding center is stable, the conformations of A1492 and A1493, which are important to binding to IF1, change drastically during the MD simulation as mentioned previously.

### 30S-IF1 conserved contacts

Multiple sequence alignment for IF1 was performed by using *ClustalW* algorithm in *JALVIEW* (version 2.11.2.7) to detect conserved residues in bacterial, archaeal and eukaryotic homologues (Clamp et al., 2004) (Extended Data Fig. 4). The most conserved residues on IF1 are located in the loop of the OB-fold region which is in close proximity to h44. Specifically, the residues Pro18, Arg23, Arg41 and Arg46 which interact with h44 (Extended Data Fig. 5) ; the residues Lys2 and Gly38 which interact with 530 loop (Extended Data Fig. 6); and Tyr60, Asp61 which interact with uS12 protein were seen to be conserved between homologues (Extended Data Fig. 7) (Carter et al., 2001). This conservation through evolution in all three kingdoms of life indicates the key role of these residues during the IF1 steering and binding for the functioning of 30S.

Our 30S^TR-SFX^ intermediate structure revealed the choreography of the initial interactions established between IF1 and 30S (Fig. 2b) (Fig. 4). We observed partially occupied electron density maps where the interactions formed with h44, uS12 and 530 loop (Extended Data Figs. 4,5,6,7), indicating the initiation of molecular interactions of IF1 near the decoding center. A loop of IF1 is introduced into the minor groove of h44, establishing connections with the nucleotide backbone, ultimately resulting in the flipping out of both A1492 and A1493. While A1493 is concealed within a pocket located on the surface of IF1, A1492 takes its position within a cavity created at the junction of IF1 and uS12 (Carter et al., 2001; DeMirci et al., 2013; Laursen et al., 2005).

Investigating the conserved interactions of IF1-uS12 protein, IF1-h44, and IF1-530 loop in the one microsecond MD simulation, the average displacements from the initial position for seven interactions of IF1-h44, i.e., Ala16—G1494, Pro18—G1494, Ala20—A1493, Arg23—G1410, Arg41—A1492, Arg46—A1493, Arg64—C1411, five interactions of IF1-uS12 protein, i.e., Ala1—Glu73, Glu3—Gly74, Tyr60—His75, Asp61—Arg41, Arg64—Leu52, and nine interactions of IF1-530 loop, i.e., Lys2—G517, Lys2—C519, Tyr35—C519, Gly38—C518, Lys39—C518, Lys39—U531, Met42—Gr530, Arg66—C519, respectively, were calculated as shown in Extended Data Fig. 8. The interfacial interactions of IF1-uS12 protein and IF1-h44 changed significantly during the dissociation, especially around 300 ns where the COM distances of IF1-uS12 protein and IF1-h44 were large. On the other hand, the interface interaction of IF1-530 loop did not change. In addition, the average displacements suggest that IF1 first dissociates from uS12 protein, and then the interaction between IF1 and h44 is loosened during the dissociation (Extended Data Fig. 8). . Thus, h44 and uS12 protein may play a role in steering IF1 deeper into the decoding center. This study represents a valuable first instance of protein-protein and protein-RNA interaction where the computational counterpart is effectively colligated with an experimental TR-SFX analysis. Notably, the observed conformation of the decoding residues validates the results obtained from MD simulations.

## Conclusion

Our time-resolved structural data have shown computationally unpredicted transient and reversible perturbations in RNA-RNA interactions (Figs. 3,5,6). Particularly disruptions in hydrogen bonding within the h44 base pairs and significant atomic distance along the P-P backbone can be attributed to the initiation of dynamic motions within the h44 region, accompanied by coordinated movements in the head and shoulder domains of the 30S. The combination of analytical methodologies with time-resolved X-ray crystallography allowed us to elucidate the *hitherto* undiscovered intricacies and unveiled novel details of early times of IF1’s binding interactions with decoding region of the 16S rRNA.

Chemical probing of IF1 binding showed enhanced reactivity of critical intersubunit bridge residue A1408 of h44 (Moazed et al. 1995) and later confirmed by crystallography and NMR (Carter et al., 2001; Dahlquist et al., 2000). As previous studies have proposed that IF1 together with IF3 induce a structure that is unfavorable for 50S docking, while the formation of 30S initiation complex promotes a conformational equilibrium that favors a favorable 50S docking structure (Milon et al., 2008; Qin et al., 2012). In other words, the binding of IF1 to h44 perturbs conformational changes and stabilizes the transition state of subunit interaction, facilitating the rate of both subunit dissociation and association. 30S^TR-SFX^ captures an alternative h44 conformation during IF1 binding suggesting that the factor may modulate the intersubunit bridge B2a/d near the decoding center. The binding of IF3 to the 30S is predicted to block the formation of the bridge B2a/d (Liu et al., 2016). As IF1 binds near the decoding center, the components of bridge B2a/d A1409 and A1410 move by 2.3 Å and 3.3 Å respectively.

Our work constitutes a significant expansion of structural studies and offers a tremendous opportunity to address fundamental structural dynamics questions of the ribosome. Building upon prior structural studies, our 30S^TR-SFX^ has directly identified transient intermediate and long-range allosteric conformational changes including the uncoupling of ribosomal h44 basepairing and global scale allosteric 30S domain closure from head rotation at the ∼200 ms time-point of IF1 binding. Long-range allosteric transitions are proposed to be at the heart of kinetic two step proof-reading model of the decoding reaction catalyzed by ribosomes (Ogle et al., 2001). Broadly, there are two models proposed for the decoding mechanism, the first one suggests that ribosome has to go from an open to close transition of 30S body in order to select the correct tRNA and the second model suggests that domain closure is not necessary (Demeshkina et al., 2012; Ogle et al., 2001). The molecular interactions identified in our work demonstrate the viability of time-resolved structural studies to address which model fits better for tRNA decoding.

Due to experimental constraints, only one of multiple possible intermediate states were probed in this study. Altogether, our data suggest that decoupling of the allosteric transitions is temporally possible, although in the absence of additional structural time-points, additional ribosomal ligands were not characterized. Much progress remains to be made, these observations have been made possible by multiple, convergent advances in XFELs, sample delivery methods, data analysis, and hybrid techniques. Our investigation of the 30S-IF1 interaction represents a single example within a one model system for the investigation of structural dynamics in biology, (Chapman et al., 2011; Fuller et al., 2017; Sierra et al., 2016;). Spanning megadaltons in their fully-assembled state, ribosomes are complex supramolecular machines and have served as a model system to explore RNA structure and protein-RNA interactions over many years. The mix-and-inject sample-delivery approach for microcrystals has fundamentally enabled time-resolved macromolecular crystallographic studies and enables capture of the conformational changes in short-lived transition states. Our TR-SFX mix-inject-probe binding experiment performed with IF1 and 30S confirmed the feasibility of investigating dynamic processes and promises to represent one of many such future explorations. For reference, the repetition rate during the beamtime in this work was 120 Hz. Improvements in X-ray pulse repetition rates and detector capabilities promise to enable data acquisition on the order of megahertz and thus reduce acquisition time per dataset. In turn, sample delivery may be adapted to facilitate collection of multiple, if not numerous, intermediate-state timepoints per XFEL beamtime. This is an exciting time for structural biology and TR-SFX provides a powerful tool for understanding the dynamic processes involved in ribosome function, such as initiation of translation. Beyond the application of the ribosome field, time-resolved crystallography studies promise to bring biology alive in general for better understanding the dynamics of macromolecular interactions at atomic resolution and near-physiological temperature.

## Methods

### MD simulation for Ribosome 30S and IF1 complex

We performed a microsecond-scale all-atom MD simulation using the structure obtained by TR-SFX. Protonation states were determined by Adaptive Poisson-Boltzmann Solver (APBS)-PDB2PQR which is a module solvation force library package with pH 7.5 (Baker et al., 2001). The modeled 30S-IF1 complex was solvated in TIP3P water using the solution builder in CHARMM-GUI so that the box size was approximately 27.3 nm in length, width and height. (Brooks et al., 2009; Jo et al., 2008; Lee et al., 2016, 2020). The solution was neutralized with 150 mM KCl. Finally, the system contained two Zn^+2^ and 254 Mg^+2^ cations, which were originally included in the 30S^TR-SFX^ structure, 2,491 K^+1^ and 1,726 Cl^-1^ anions, and about 600,000 water molecules in addition to 30S, thus the system became very large size, containing approximately 1.9 million atoms in total.

MD simulation was performed using *Amber* software, with the all-atom additive *CHARMM36m* force field for protein and rRNA residues since the structure has several modified nucleic residues (7MG, M2G, 5MC, 2MG, 4OC, 3MU, M6A) (Case et al., 2005; Huang et al., 2017). The simulation was performed undmiyaer periodic boundary conditions (Case et al., 2005). In the simulation, the time step was 2 fs and the trajectory interval was 10 ps. The temperature was maintained at 298.15 K (ambient temperature) using a *Langevin* thermostat, and the pressure was maintained at 1 bar using a *Berendsen* barostat. The shake method was also employed (Miyamoto & Kollman,1992; Ryckaert et al., 1977). Short-range electrostatic and van der Waals forces were cut off at 12 Å. First, energy minimization was performed with restraint to the solute and no pressure control, and energy minimization without any restraint was executed to remove bad contacts. The temperature of the system was heated to 298.15 K in 100 ps while a 10.0 kcal/(mol Å^2^) constraint was applied to the solution using a harmonic restraint. The *Langevin* thermostat was used for the system heating and an anisotropic *Berendsen* weak-coupling barostat was used to also equilibrate the pressure. After the heating process, we performed a 5 ns equilibration process with a larger skinnb value in order to equilibrate the dimensions and density of the systems. Finally, the temperature was controlled using the *Langevin* thermostat using a constant pressure periodic boundary with an average pressure of 1 atm, and we performed one microsecond MD simulation.

### Purification of 30S ribosomal subunits

30S ribosomal subunits from *T. thermophilus* HB8 (ATCC27634) were purified as described (DeMirci et al., 2010, 2013). The *T. thermophilus* HB8 strain was grown in 4 L baffled glass flasks using ATCC:697 *Thermus* medium enriched with Castenholz salts, at a temperature of 72 ◻ with intense aeration at 400 rpm by using New Brunswick Innova 4430R rotating incubator. The cells were harvested by centrifugation at 4000 rpm at 4 ◻ when their optical density reached OD_600_=0.8. The cell pellets were then cooled by flash freezing in liquid nitrogen and stored at - 80 ◻ until further use. To resuspend the cells, Buffer A containing 100 mM ammonium chloride (NH_4_Cl), 10.5 mM magnesium acetate (MgOAc), 0.5 mM Ethylenediaminetetraacetic acid (EDTA), 20 mM N-2-hydroxyethylpiperazine-N-2-ethane sulfonic acid (HEPES), adjusted with potassium hydroxide (KOH) to pH 7.5 was used. Resuspended cells were disrupted by two passes through an emulsiflex C5 (Avestin, ON, Canada) initially at 12,000 psi and a second pass at 20000 psi. After disruption, lysate was centrifuged at 20000 rpm for 20 minutes with a Ti-45 rotor (Beckman Coulter, USA) to remove the cell debris. Supernatants of the sample applied to sucrose cushion Buffer B containing 1.0 M NH_4_Cl, 1.1 M sucrose, 10.5 mM MgOAc, 0.5 mM EDTA, 20 mM HEPES-KOH (pH 7.5) were centrifuged at 43000 rpm for 15 hours with a Ti-45 rotor. After rewashing the samples with Buffer A, pellets were resuspended with Buffer C containing 1.5 M ammonium sulfate (NH_4_)_2_SO_4_, 10 mM MgOAc, 20 mM HEPES-KOH (pH 7.5) and loaded onto the POROS-ET hydrophobic reverse phase interaction column. Reverse ammonium sulfate gradient containing Buffer C and Buffer D containing 20 mM HEPES-KOH (pH 7.5) and 10 mM MgOAc was used for elution of samples, and 70S peak fractions were collected. The collected fractions were pooled and dialyzed into Buffer E containing 50 mM KCl, 10 mM NH_4_Cl, 10 mM HEPES-KOH (pH 7.5), 10.25 mM MgOAc, 0.25 mM EDTA. Separation of the 30S and 50S subunit fractions was achieved by applying a 10% - 40% linear sucrose gradient in Buffer F containing 50 mM KCl, 10 mM NH_4_Cl, 10 mM HEPES-KOH (pH 7.5), 2.25 mM MgOAc, 0.25 mM EDTA. The 30S peak was then dialyzed into Buffer C for further separation by the POROS-ET column with a reverse ammonium sulfate gradient. The RNase-free fractions were dialyzed into Buffer G containing 50 mM KCl, 10 mM NH_4_Cl, 5 mM HEPES-KOH (pH 7.5), 10 mM MgOAc and concentrated to the final OD_260_=400 using Amicon 30 KDa PM30, an ultrafiltration membrane.

### Crystallization of 30S ribosomal subunits

The purified 30S^APO^ was crystallized by the hanging drop method at 4 °C with the 1:1 mixture containing buffer G and the reservoir solution buffer H (16% (v/v) 2-methyl-2,4-pentanediol (MPD) in buffer 15 mM MgCl_2_, 100 mM KCl, 50 mM NH_4_Cl, 100 mM 2-(N-morpholino) ethanesulfonic acid MES buffer (pH=6.5)). The 3–10 × 3–10 × 20–30 μm^3^ sized microcrystals were harvested in the buffer G to be pooled. Crystal density was approximated to be 10^9^–10^10^ particles per ml based on light microscopy and nanoparticle tracking analysis NanoSight LM10-HS with corresponding Nanoparticle Tracking Analysis (NTA) software suite (Malvern Instruments, Malvern, UK).

### CoMESH construction and operation

The concentric microfluidic electrokinetic sample holder (coMESH) comprises an outer-flow capillary and an inner-flow capillary, allowing simultaneous liquid injection (Sierra et al., 2016). The lengths of these capillaries are adjustable, providing control over the mixing time before introducing the liquids into the vacuum chamber at the interaction point (Fig. 1a). The inner capillary (100◻μm ID and 160◻μm OD) is filled with the 30S^APO^ crystal slurry in its native *mother liquor,* while the outer capillary (200◻μm ID and 360◻μm OD) contains 10 mg/ml IF1 in a *sister liquor* to be mixed with the 30S^APO^ sample to enable time-resolved data collection during injection (Sierra et al., 2016). Diffusion coeeficient of the IF1 is calculated as 13.214 ◻^2^/ns using the following equation, 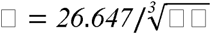 (Xie et al., 2017). Here, MW represents the molecular weight of the IF1, which is 8.2 kDa. The diffusion times are calculated as 2.924 ms and 8.519 ms for crystal sizes 3μm⊡3μm⊡5μm and 5μm⊡5μm⊡10μm, respectively. This calculation employs the following equation:

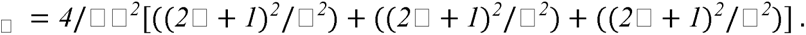

Here ‘a’,’b’ and ‘c’ represent the half-length of the crystal, and we set ‘l’, ‘m’, and ‘n’ to 0 for the calculation of the slowest diffusion rate _◻_.

To determine the time dependant mixing concentration, we employed the following equation:

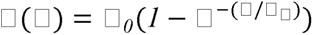 (Schmidt, 2013). This equation corresponds to the mixing time range between 80 ms and 240 ms. Prior to connecting the 30S^APO^ sample injection line, the IF1 carrying sister liquor is loaded, flowed and subjected to electrical focusing. The 30S^APO^ sample injection line is connected when the liquid-jet is nearly stabilized. It is worth noting that the sister liquor never reaches full stabilization, as reported in the literature (Sierra et al., 2016). Also, all flows mentioned here are considered as laminar flows. The flow rates of the 30S^APO^ sample line and sister liquor carrying IF1 are nearly identical to ensure a stable injection and to maximize the number of crystals exposed to X-ray at the interaction point. The flow rate of the sample line is gradually increased until diffraction is detected.

### Sample injection

The concentrically mixed 30S^APO^ microcrystals by electrospinning for ∼200 ms duration were transported to the X-ray interaction region using the coMESH system (Sierra et al., 2016) with a 1◻μl/min nominal flow rate for each syringe pump (Fig. 1a). To study the time-resolved 30S-IF1 complex, we employed the coMESH system for efficient sample injection. The sample reservoir was loaded with a slurry of unmixed 30S ribosome microcrystals in the original mother liquor. The outer sister liquor, devoid of 30S microcrystals, and containing IF1 protein solution with the same buffer as the mother liquor, with an increased MPD (2-methyl-2,4-pentanediol) concentration of 20% (v/v) to aid in vacuum injection (Sierra et al., 2016). The 30S^APO^ ribosomal subunit microcrystals were in the inner sample line, suspended in their native mother liquor containing 16% (v/v) MPD. These microcrystals displayed a uniform size distribution, ranging from 3-5μm⊡3-5μm⊡5-10μm in each dimension due to controlled slower growth at a temperature of 4 °C. Larger crystals were selectively removed through repeated gentle differential settling, avoiding centrifugation (Sierra et al., 2016). The injection system involved three capillaries: an inner sample capillary made of fused silica, an outer sheath flow capillary made of fused silica, and an outer concentric capillary tapered. The tips of the inner and outer capillaries were positioned coterminally at the exact location (Sierra et al., 2016). During the injection process, a voltage of between 3-4 kV to maximize data collection was applied to the sheath liquid, while the counter electrode remained grounded potential. The sample flowed at a rate of 1 μl/min, and the sheath flow rate was adjusted to match the sample flow rate at 1 μl/min. For a given capillary size, this corresponds to a flow rate of 3.32 x 10^-11^ m^3^/s. (Sierra et al., 2016).

### Data collection and analysis

The SFX data collection was conducted at the CXI instrument within the Linac Coherent Light Source (beamtime ID: cxils9717 during LCLS run 17), at the SLAC National Accelerator Laboratory in Menlo Park, California, under controlled ambient temperature conditions. An X-ray beam, characterized by vertically polarized and the pulses lasting about 40 femtoseconds, was focused employing refractive beryllium compound lenses, resulting in an approximate beam size of 6 × 6 µm² at full width at half maximum (FWHM). The experimental procedures were executed employing a photon energy of 9.5 keV, and the data collection operated at a repetition rate of 120 Hz utilizing SASE mode with a wavelength of 1.29 Å at a 293 K. Real-time data analysis was systematically conducted to establish the initial diffraction geometry, monitor crystal hit rates, and analysis of the gain-switching modalities of the CSPAD-2.3M detector (Kameshima et al., 2014). This analytical endeavor was facilitated through the *OM monitor* version 1.0 (Mariani et al., 2016) and *Psocake* version 1.0.8 (Damiani et al., 2016; Thayer et al., 2017; Yoon, 2020). Data collected from *T. thermophilus* 30S microcrystals continuously during 216 minutes and 58 seconds of beamtime. A total of 1,540,800 detector frames were collected. These *T. thermophilus* 30S microcrystals were delivered to the X-ray interaction point, using the coMESH system (Sierra et al., 2016). This method yielded a total of 148,000 registered hits. Each individual diffraction pattern hit was defined as frames displaying more than 30 discernible Bragg peaks, each exhibiting a minimum signal-to-noise ratio of not less than 4.5.

### Structure determination and refinement

The intermediate state structure of 30S^TR-SFX^, which captures the transient binding of IF1 to 30S, was determined at 3.59 Å resolution in space group P4_1_2_1_2. Molecular replacement was performed using the *PHASER* program (McCoy et al., 2007) within the *PHENIX* version 1.19.2 software suite (Adams et al., 2010). Previously published 30S structure (PDB ID: 4DR1) was used as the initial search model (DeMirci et al., 2013). Initial rigid body refinement was performed with the coordinates of the 30S ribosomal subunit (PDB ID: 4DR1) by using *PHENIX* version 1.19.2 software suite (Adams et al., 2010). Subsequently, IF1 was manually copied and reconstructed to the 30S ribosomal subunit in *COOT*, guided by the 30S-IF1 complex structure (PDB ID: 1HR0)(Carter et al.,2001). Further structure refinement was performed by using individual coordinates and TLS parameters, following the simulated-annealing refinement. Potential positions of altered side chains and water molecules were identified by performing composite omit map refinement within *PHENIX*. The electron density map was verified in *COOT* version 0.8.9.2 (Emsley & Cowtan, 2004). During this process, the altered positions with strong difference density were retained and hexahydrated magnesium molecules that were located outside the electron density were manually removed. The Ramachandran statistics for the 30S^TR-SFX^ structure are as follows: favored / allowed / outliers = 85.9 /13.9 /0.1 %, respectively. The detailed structure refinement statistics are indicated in the Extended Data Table 1. For the representation of structure, alignments and figure generation, *PyMOL* version 2.3 (DeLano, 2002) was used and multiple sequence alignment was performed by using *JALVIEW* 2.11.2.7 (Clamp et al., 2004).

## Supporting information

https://docs.google.com/document/d/16mcoXvx-_T5FuEnXb9dowLt2cobZQIn0y9o3QpBvCKE/edit?usp=sharing

## Data availability

Coordinates and structure factors of the 30S^TR-SFX^ structure have been deposited at the Protein Data Bank (PDB) under accession code 8WRC (Time-Resolved Ambient Temperature Kineto-Crystallographic Structure of Initiation Factor in Complex with Ribosome).

## Code availability

Conducted a comparison of the distances between the phosphate atoms of the 30S ribosomal subunit in both its IF1-bound and unbound conformation. Utilized ’*atom-wise distances between matching AtomGroups*,’ which allowed us to compute the distances between atom groups containing the same number of atoms. The code for pairwise P-P distance calculations is accessible through MDAnalysis 0.19.0; https://doi.org/10.1002/jcc.21787, and three-dimensional pairwise visualization was facilitated using MDAnalysis in conjunction with *PyMOL*.

## Acknowledgements

Authors would like to dedicate this manuscript to the memory of Dr. Albert E. Dahlberg and Dr. Nizar Turker. We thank the staff at Linear Coherent Light Source at SLAC National Accelerator Laboratory for their assistance in data collection. We thank the Photon Ultrafast Laser Science and Engineering Institute (PULSE Institute) at Stanford University for providing technical support. The numerical calculations were conducted in part using Cygnus and Pegasus at the Center for Computational Sciences, University of Tsukuba.

## Author Contributions

The project was initiated and coordinated by H.D.; H.D., Y.R., B. H., and E.H.D, supported crystallography applications at LCLS.; In a beamtime M.H., M.L., C.K., R.G.S. prepared the X-ray instrument and collected the data.; Initial ribosome crystals were developed in an experiment with H.D., E.H.D.; Highly diffracting crystals were optimized and tested for diffraction at the SSRL BL12-2 by H.D.; Ribosomes are purified and crystallized by H.D.; The coMESH sample injection was optimized by R.G.S., H.D., and E.H.D.; Sample preparation and reservoir loading at the LCLS was performed by H.D., C.K and R.G.S.; coMESH injectors operated during the XFEL beamtimes by R.G.S., and E.H.D.; Data processing during the beamtime was performed by F.P.; with the help of C.Y.; E.H.D., Y.R., and H.D recorded progress during data collection.; Data processing was further completed by F.P.; Structures were refined by I.Y.;, and data were interpreted by I.Y. with the help of E.H.D., S.Y., E.D., E.A., A.S., FBE, CKul, M.Y., B.T., H.I.C., A.K., J.J., O.G., A.E., P.M., A.M., S.W., and H.D.; Molecular dynamics simulations were analysed by S.Y. and A.M.; 2D and 3D pairwise distance analysis was performed by E.A.; The TR-SFX experiment was designed by H.D. and R.G.S.; All of the authors read and acknowledged the manuscript.

## Additional information

### Supplementary Information

Supplementary Information is available for this paper.

**<u>Supplementary Video 1</u>. Morph representation of the motion of critical residues A1492 and A1493 in the decoding center of 30S.** The movie is generated by using Morph feature of *PyMOL*. For the visualizing the flip-out and flip-in conformation of the residues A1492 and A1493 at the decoding center in the presence of IF1 binding, first two structure is aligned: 30S^APO^ and 30S^TR-SFX^; further 30S^TR-SFX^ and 30S^HOLO^ structures aligned, respectively. Two morph are generated as result of 2 structure alignments by using mporh command in the *PyMOL*. The movie is created by the combination of two morphs in *PyMOL*.

**<u>Supplementary Video 2</u>. Conformational changes of A1492 and A1493 in the one microsecond MD simulation.** The MD trajectories were visualized on visual molecular dynamics (VMD). The drawing method and color scheme, respectively, of each domain is as follows: IF1, new cartoon, blue; A1492 and A1493, licorice, pink. A1492 and A1493 are the lower and upper nucleic acid residues, respectively. The conformations of A1492 and A1493 changed significantly. Around 300 ns, the side chain of A1492 transitioned into the tucked-in conformation, resembling the 30S^APO^ state. After this A1492 conformational change, the structural fluctuations of A1493 increased, resulting in the A1493 side chain also swinging away from IF1.

**<u>Supplementary Video 3</u>. Conformational changes of A1492 and A1493 in the additional 100 ns MD simulation.** The drawing method and color scheme, respectively, of each domain on VMD is as follows: IF1, new cartoon, blue; A1492 and A1493, licorice, pink. A1492 and A1493 are the lower and upper nucleic acid residues, respectively. The additional 100 ns MD simulation using the structure at 1 μs as the initial structure showed that A1493 formed the fully tucked-in conformation, which is similar to the conformation in 30S^APO^ state.

**<u>Supplementary Video 4</u>. Dynamics of decoding center in the MD simulation of the 30S-IF1 complex.** The drawing method and color scheme, respectively, of each domain on VMD is as follows: IF1, surf, blue; r44, surf, pink; uS12 protein, surf, green; 530 loop, surf, yellow; A1492, licorice, cyan; A1493, licorice, orange. At 0 ns, IF1, uS12 protein, and h44 were tightly packing, and the interactions between them were strong. However, their interactions were loosened by 300 ns, and IF1 dissociated slightly from the decoding center. On the other hand, the interactions between IF1 and 530 loop was stable.

**<u>Supplementary Video 5</u>., <u>Supplementary Video 6</u>. Perturbations of the base interactions within G1423-U1427 and A1473-C1477 in the MD simulation.** In the videos, the Apo-state and Holo-state structures are shown in yellow and cyan, respectively, as reference structures for comparison, and the MD simulation results are shown in pink. The left and right structures are G1423-U1427 and A1473-C1477, respectively, and MD simulations were superimposed on all atoms except hydrogen atoms in A1473-C1477 of the reference structures. The perturbations of the base interactions within h44 were observed, and the conformations in the MD simulation were similar to the 30S^APO^ state.

**<u>Supplementary Video 7</u>.**, **<u>Supplementary Video 8</u>. Perturbations of the base interactions within C1412-G1415 and U1485-G1488 in the MD simulation.** In the videos, the Apo-state and Holo-state structures are shown in yellow and cyan, respectively, as reference structures for comparison, and the MD simulation results are presented in pink. The left and right structures are C1412-G1415 and U1485-G1488, respectively, and MD simulations were superimposed on all atoms except hydrogen atoms in U1485-G1488 of the reference structures. The perturbations of the base interactions within h44 were observed, and the conformations in the MD simulation were similar to the 30S^APO^ state.

**<u>Supplementary Video 9</u>. The analysis involved the assessment of 3D pairwise distance motion within the 30S^APO^, 30S^TR-SFX^, and 30S^HOLO^ structures.** A comparative examination was carried out to assess the variations in the distances between phosphate atoms within the 30S ribosomal subunit in both its IF1-bound and unbound states. The dynamic visual representation was generated using PyMOL by implementing MDAnalysis source codes. The alignment of the 30S^APO^ structure with the 30S^TR-SFX^ structure revealed a remarkable maximal spatial shift of 6 Å, particularly within the residues A1408-G1422, consequently resulting in significant h44 motions relative to the original 30S^APO^ structure. The initial binding of IF1 induced deliberate lateral shifts of approximately 2.0 Å, particularly in the region of residues G75-U98 and A463-U480. A detailed investigation into the three-dimensional pairwise distances between the 30S^TR-SFX^ and 30S^HOLO^ structures disclosed subtle displacements, ranging from 2 to 3 Å, among residues A453-G485, G144-U204, A665-G710, and G75-U98. Upon superimposing the 30S^TR-SFX^ and 30S^HOLO^ structures, the intermediate binding of IF1 to the 30S structure initiated h44 motions and head movements spanning a range of 2.0 to 4.0 Å, primarily affecting residues between A938-G1454.

**<u>Supplementary Video 10</u>., <u>Supplementary Video 11</u>. The structural fluctuations from the average structure by principal component analysis.** Principal component analysis was performed on the phosphate atoms of rRNA in the one microsecond MD simulation. We visualized the structural fluctuation from the average structure of each phosphate atom along the first principal component (PC1). Supplementary Videos 10 and 11 show the structural fluctuations of rRNA from a top down view and from a head view, respectively. The magnitude of the original fluctuation was shown by five times. The large-scale movements of 30S in the MD simulation had similar structural fluctuation tendency as those in the 16S rRNA pairwise comparative analysis of 30S^TR-SFX^ and 30S^HOLO^ except for C150-G168, G674-G714, C1254-C1284 and the decoding center of h44. Also, C1254-C1284, which showed large fluctuations, is shown in red, h44 is shown in pink, the others in A938-G1454 are colored in blue, A1492 and A1493 are shown in green, and the others that do not correspond to them are shown in white.

