## Supplementary material for "4D Crystallography Captures Transient IF1-Ribosome Dynamics in Translation Initiation": https://docs.google.com/document/d/16mcoXvx-_T5FuEnXb9dowLt2cobZQIn0y9o3QpBvCKE/edit?usp=sharing

**Extended Data Table 1 Data collection and refinement statistics for 30S^TR-SFX^ structure at ~200 ms time point**

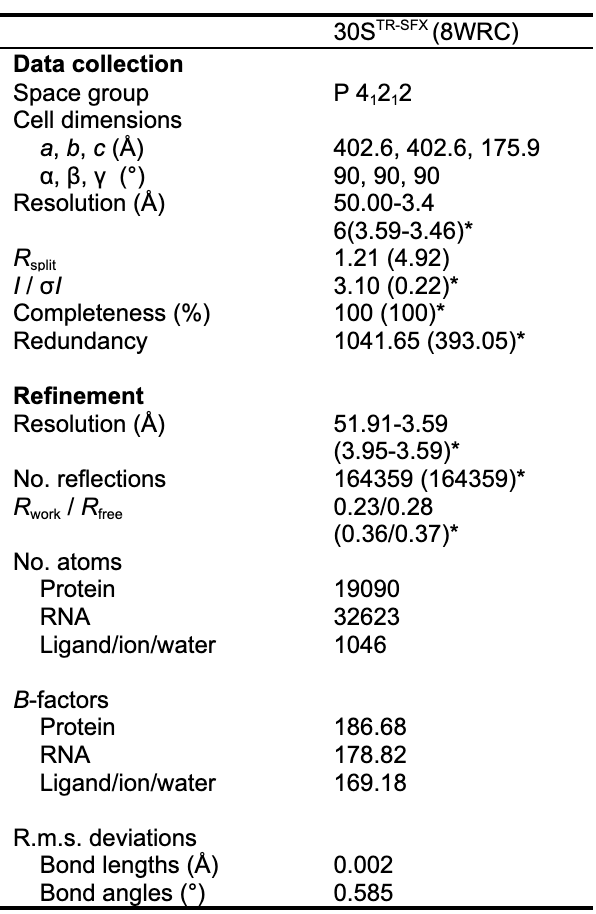

*150,000 crystals used for the dataset.

*Values in parentheses are for highest-resolution shell.

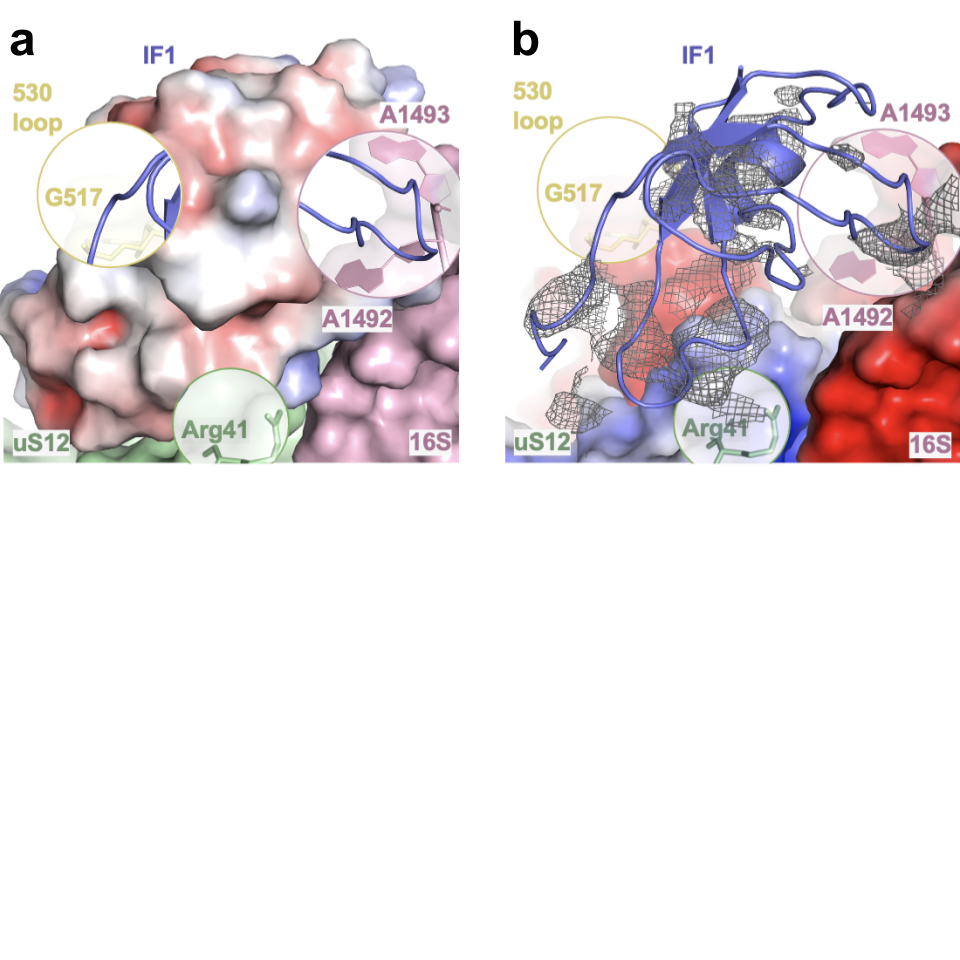

**Extended Data Fig. 1| Electrostatic representation of IF1 binding region. a,** IF1 and **b,** its surroundings (16S rRNA, S12 and 530 loop) are represented with electrostatic to indicate the charge forces during the the binding of IF1 to 30S. APBS electrostatics are generated in *PyMOL*.

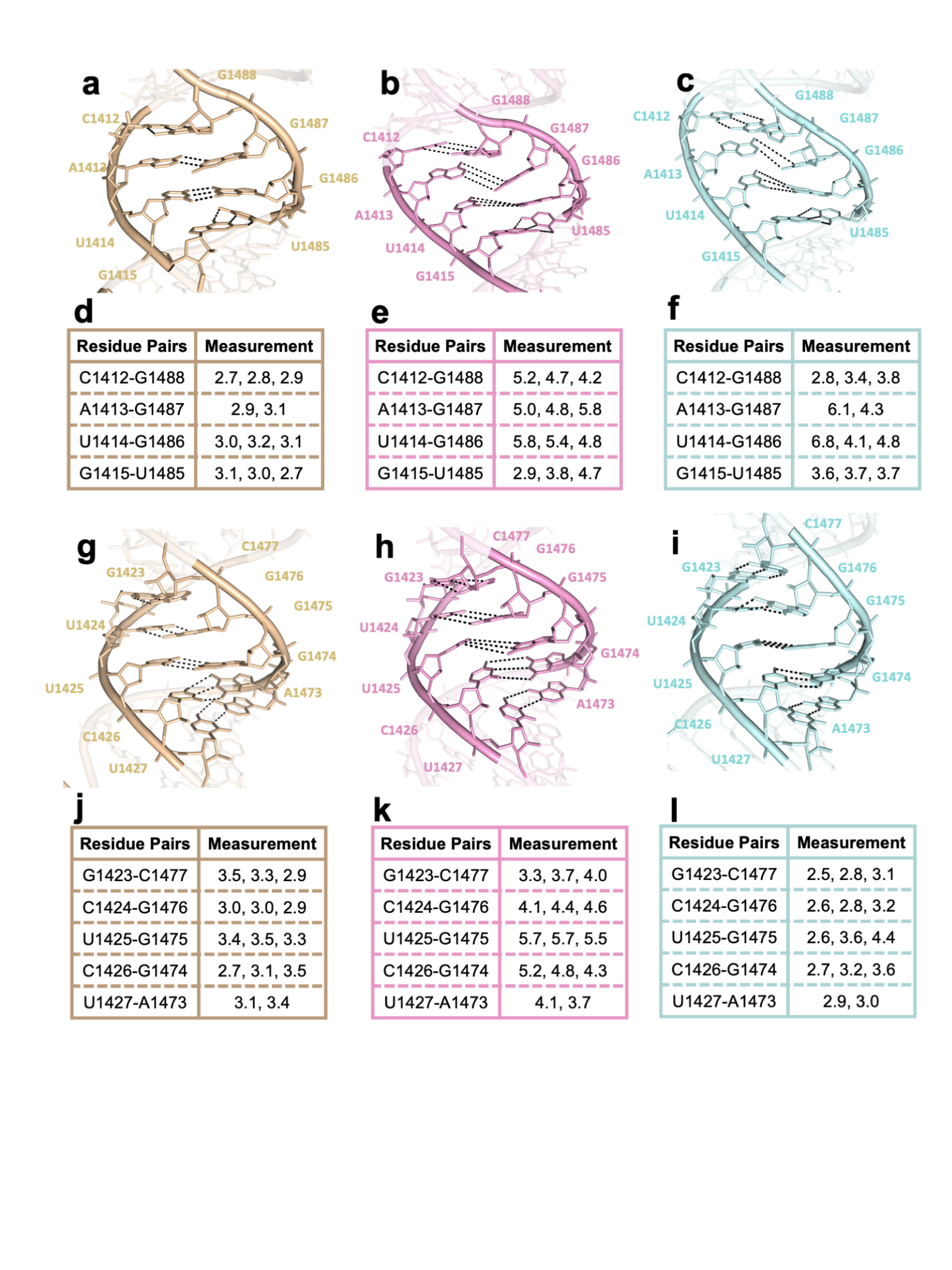

**Extended Data Fig. 2 | Measurement of Distances between Base Interactions in the h44 Region of the 16S rRNA Structure during IF1 Binding. a, b, c,** Depict the perturbations of base interactions in the 30S^APO^, 30S^TR-SFX^, and 30S^HOLO^ h44 structures respectively, illustrating the distances between the residue pairs C1412-G1488, A1413-G1487, U1414-G1486, G1415-U1485, respectively. **d, e, f,** This figure comprises tables that quantify the distances between the specified base pairs for these structures. **g, h, i,** Show the alterations in base interactions in the 30S^APO^, 30S^TR-SFX^, and 30S^HOLO^ h44 structures respectively, demonstrating the distances between the residue pairs G1423-C1477, C1424-G1476, U1425-G1475, C1426-G1474, and U1427-A1473, respectively. **j, k, l,** Similarly, this figure contains tables that quantify the distances between the specified base pairs for these structures.

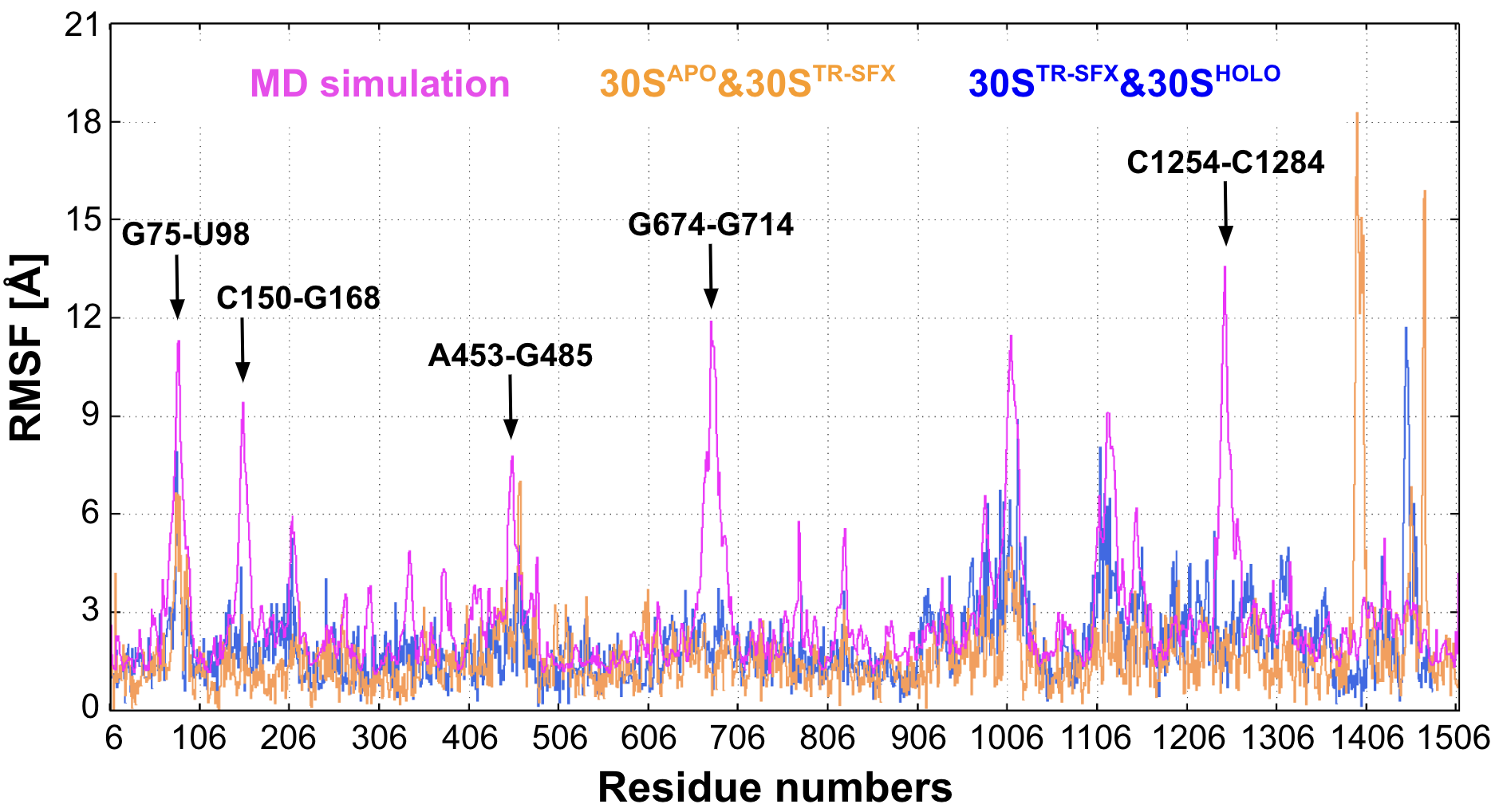

**Extended Data Fig. 3. Comparison between RMSF for the MD simulation of 30S-IF1 complex and two-dimensional (2D) pairwise distances between the 30S^APO^ and 30S^TR-SFX^ structures and between the 30S^TR-SFX^ and 30S^HOLO^ structures.** The RMSF, 2D pairwise distance between the 30S^APO^ and 30S^TR-SFX^ structures, and 2D pairwise distance between the 30S^TR-SFX^ and 30S^HOLO^ structures correspond to pink, orange, and blue, respectively. The RMSF for only the phosphate atoms in the MD simulation was computed for comparison with the two-dimensional (2D) pairwise distances. The two-dimensional (2D) pairwise distances were multiplied by three to compare the trend with the RMSF. In the MD simulation, there are large structural fluctuations around G75-U98, C150-G168, A453-G485, G674-G714, and A938-G1454 (specifically, C1254-C1284), which correspond to large movements as results of principal component analysis. The large-scale movements of 30S in the MD simulation had similar structural fluctuation tendency as those in the 2D pairwise comparative analysis of 30S^TR-SFX^ and 30S^HOLO^ except for C150-G168, G674-G714, C1254-C1284 and the decoding center of h44.

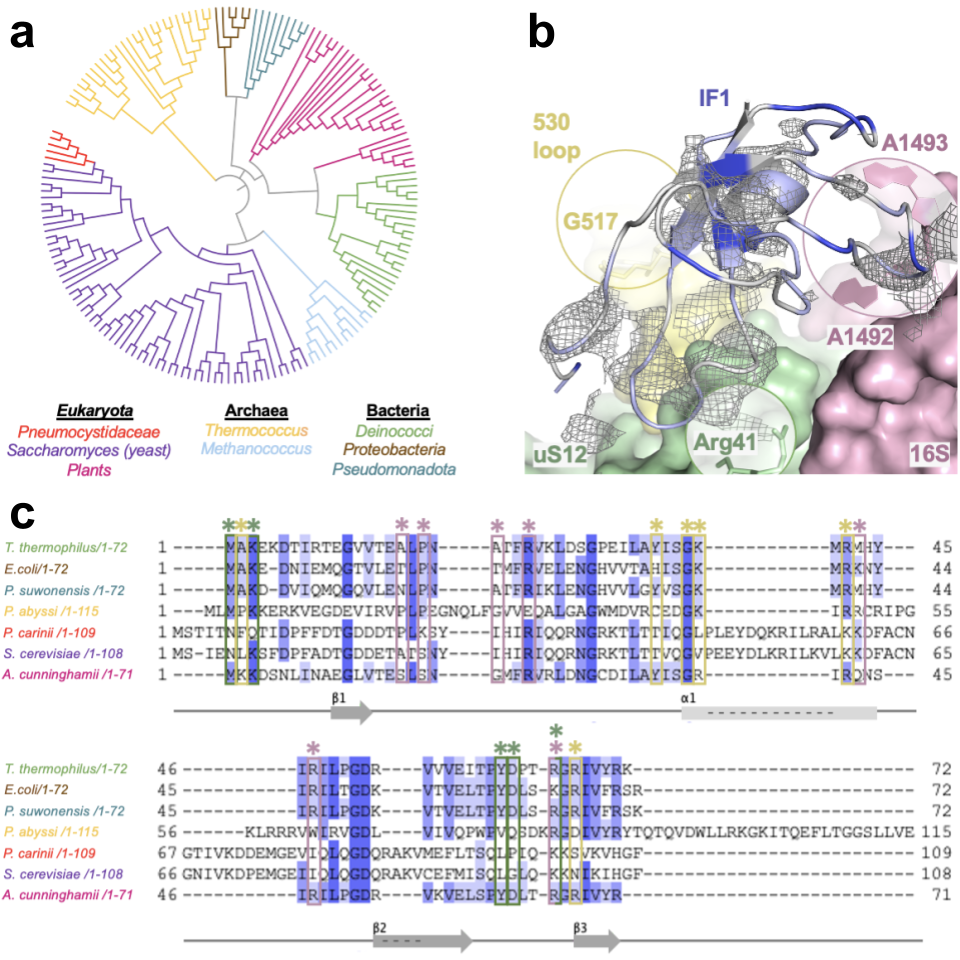

**Extended Data Fig. 4.** Multiple sequence alignment based on homologues of IF1. **a,** Phylogenetic tree is builded to represent the evolutionary relationships among homologues of IF1. **b,** Conserved residues are highlighted on the IF1 of 30S. **c,** Multiple sequence alignment is performed by using *ClustalW* in *Jalview 2.11.2.7*. Bacterial sequences are colored based on percentage identity and interacting residues of IF1 with 16S rRNA, S12 protein and 530 loop are indicated with pink, green and yellow stars/squares, respectively.

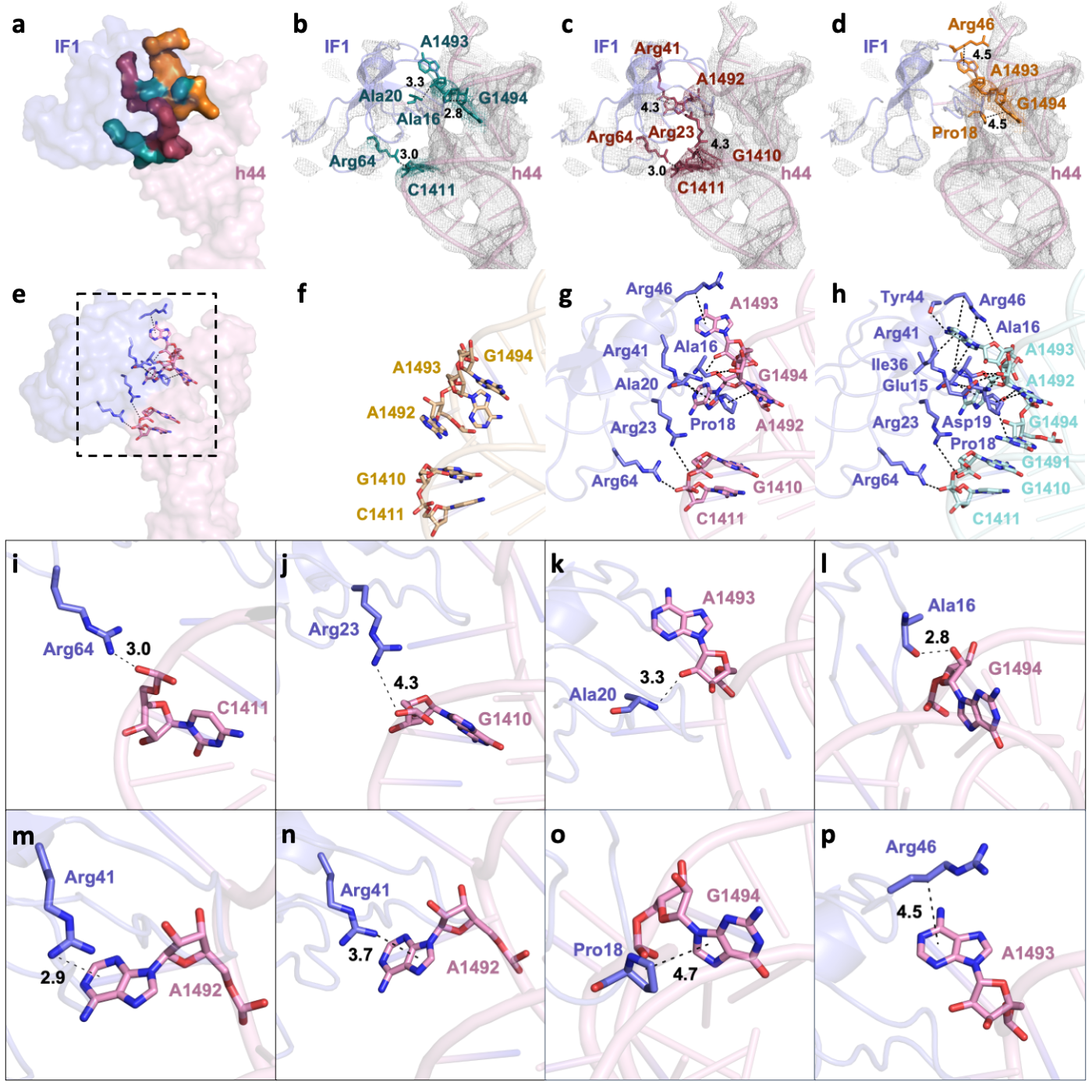

**Extended Data Fig. 5. Possible interactions at the interface between IF1 and h44 in 30S^TR-SFX^ structure. a,** Surface representation of interaction site in IF1 (slate) and h44 (pink), with possible hydrogen bond, electrostatic interactions and hydrophobic interactions are highlighted in deepteal, raspberry and tv-orange colors, respectively. **b,c,d,** Stick representation of possibly interacting residues with hydrogen-bond (deep teal), electrostatic (raspberry) and hydrophobic (tv-orange) in their *2Fo-Fc* electron density maps contoured at 1σ level. Localization of h44 interacting residues with IF1 is shown in **f,** 30S^APO^ **g,** 30S^TR-SFX^ and **h,** 30S^HOLO^ structures. h44 is colored in wheat, pink and cyan respectively, while IF1 is represented in slate. The possible interacting residues in their stick representations shown together in **e,** and individually in **i, j, k, l, m, n, o** and **p.**

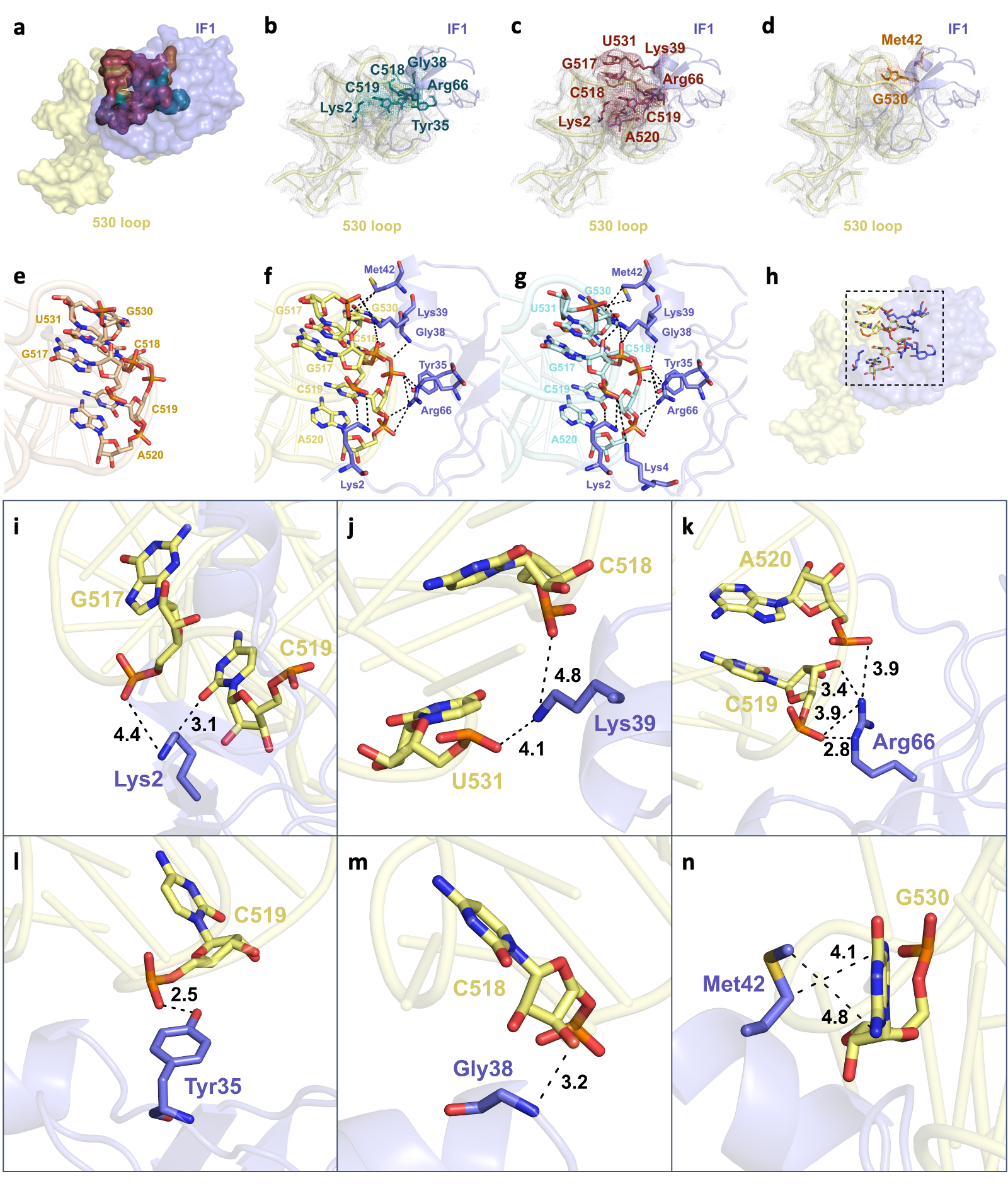

**Extended Data Fig. 6. Possible interactions at the interface between IF1 and 530 loop in 30S^TR-SFX^ structure. a,** Surface representation of interaction site in IF1 (slate) and 530 loop (yellow), with possible hydrogen bond, electrostatic interactions and hydrophobic interactions are highlighted in deepteal, raspberry and tv-orange colors, respectively. **b,c,d,** Stick representation of possibly interacting residues with hydrogen-bond (deep teal), electrostatic (raspberry) and hydrophobic (tv-orange) in their *2Fo-Fc* electron density maps contoured at 1σ level. Localization of 530 loop interacting residues with IF1 is shown in **e,** 30S^APO^ **f,** 30S^TR-SFX^ and **g,** 30S^HOLO^ structures. 530 loop is colored in wheat, yellow and cyan respectively, while IF1 is represented in slate. The possible interacting residues in their stick representations shown together in **h,** and individually in **i, j, k, l, m, n.**

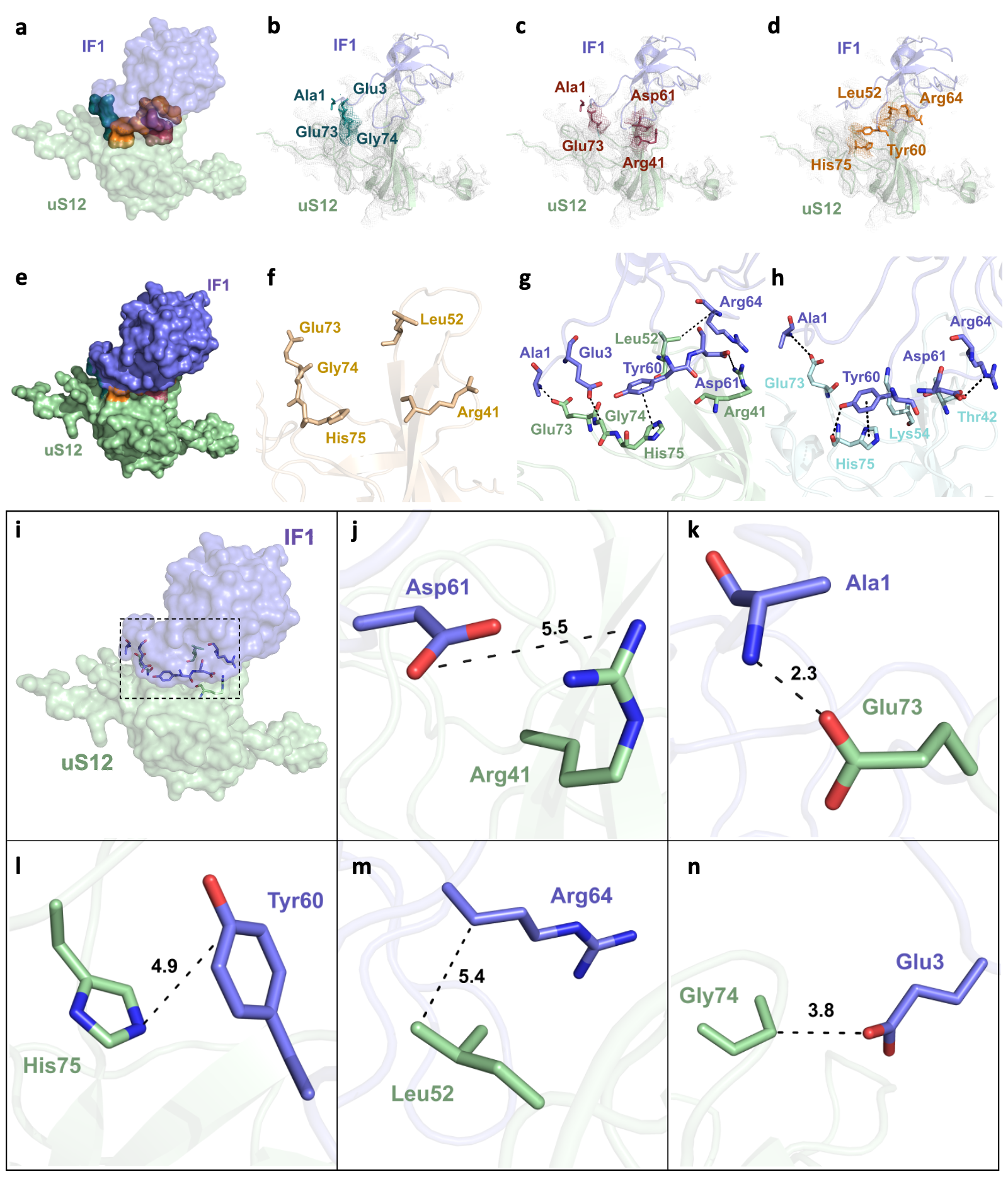

**Extended Data Fig. 7. Possible interactions at the interface between IF1 and uS12 protein in 30S^TR-SFX^ structure. a,e** Surface representation of interaction site in IF1 (slate) and uS12 protein (green), with possible hydrogen bond, electrostatic interactions and hydrophobic interactions are highlighted in deepteal, raspberry and tv-orange colors, respectively. **b,c,d,** Stick representation of possibly interacting residues with hydrogen-bond (deep teal), electrostatic (raspberry) and hydrophobic (tv-orange) in their *2Fo-Fc* electron density maps contoured at 1σ level. Localization of uS12 interacting residues with IF1 is shown in **f,** 30S^APO^ **g,** 30S^TR-SFX^ and **h,** 30S^HOLO^ structures. uS12 is colored in wheat, yellow and cyan respectively, while IF1 is represented in slate. The possible interacting residues in their stick representations shown together in **i,** and individually in **j, k, l, m, n.**

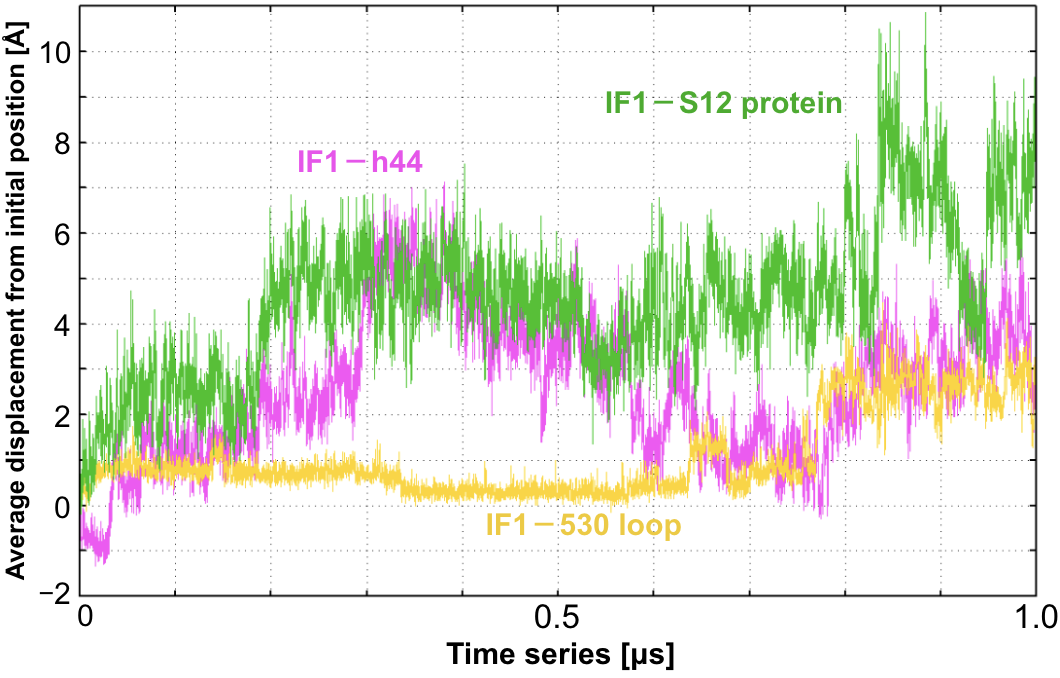

**Extended Data Fig. 8. Time series of average displacements from the initial position for 30S-IF1 conserved contacts.** The average displacements from the initial structure were calculated for seven interactions of IF1ｰh44, i.e. Ala16ｰG1494, Pro18ｰG1494, Ala20ｰA1493, Arg23ｰG1410, Arg41ｰA1492, Arg46ｰA1493, Arg64ｰC1411, for five interactions of IF1ｰuS12 protein, i.e. Ala1ｰGlu73, Glu3ｰGly74, Tyr60ｰHis75, Asp61ｰArg41, Arg64ｰLeu52, and for nine interactions of IF1ｰ530 loop, i.e. Lys2ｰG517, Lys2ｰC519, Tyr35ｰC519, Gly38ｰC518, Lys39ｰC518, Lys39ｰU531, Met42ｰGr530, Arg66ｰC519, Arg66ｰA520, respectively. First, the interactions between IF1 and uS12 protein were loosened. When the average displacement between IF1 and uS12 protein was about 2 Å away, the interaction between IF1 and h44 became unstable, and IF1 started dissociating from h44 as well. Once IF1 dissociated from uS12 protein and h44, their interfacial interactions changed significantly. On the other hand, the interactions between IF1 and the 530 loop were relatively stable.
